# Systematic functional analysis of the Com pilus in *Streptococcus sanguinis*: a minimalistic type 4 filament dedicated to DNA uptake in monoderm bacteria

**DOI:** 10.1101/2023.09.19.558370

**Authors:** Jeremy Mom, Iman Chouikha, Odile Valette, Laetitia Pieulle, Vladimir Pelicic

## Abstract

Type 4 filaments (T4F) are a superfamily of functionally versatile nanomachines, ubiquitous in prokaryotes, which use similar multi-protein machineries to assemble and operate filamentous polymers of type 4 pilins. The best studied T4F use very complex machineries, which has posed challenges to understanding the mechanisms of both filament assembly and the roles they facilitate. Here, we report the systematic functional analysis of the Com pilus, a widespread T4F mediating DNA uptake during natural transformation in monoderm bacteria. Using *Streptococcus sanguinis* as a model, we show that Com pili are *bona fide* type 4 pili (T4P), which represent a new pilus sub-type. We show that with only eight components necessary for their assembly and functioning – all “core” poteins universally conserved across this superfamily – the Com pilus epitomises a minimalistic T4F. We demonstrate that core T4F components are sufficient for filament assembly. Intriguingly, akin to more elaborate T4F, the Com pilus contains four minor pilins forming a complex likely to be situated at the apex of the filaments. Our results have global implications for T4F and make Com pili a model for elucidating the fundamental processes underpinning filament assembly.

## Introduction

Type 4 filaments (T4F) are a superfamily of nanomachines found ubiquitously in Bacteria and Archaea [1], which mediate an astonishing diversity of functions including adhesion, motility, protein secretion, DNA uptake *etc*. T4F use a conserved multi-protein machinery assembling and operating a filamentous polymer of type 4 pilins [2]. How these filaments are assembled and mediate such different functions remain poorly understood.

The best characterised T4F are widespread virulence factors in diderm bacterial pathogens: type 4a pili (T4aP, where “a” denotes the sub-type) and type 2 secretion systems (T2SS). Although they produce very different filaments – µm-long pili readily detectable on the cell surface (T4aP), or elusive endopili (also known as pseudopili) confined to the periplasm (T2SS) – these T4F use almost identical multi-protein machineries [3]. The ∼ 15 conserved components form a multi-layered structure spanning the entire cell envelope from cytoplasm to outer membrane [4]. However, only four core components are universally conserved in T4F and are therefore suspected to play a role in filament assembly [3]: type 4 pilins (usually one major and several minor pilins, often poorly characterised), prepilin peptidase (PPase), extension ATPase and platform protein. Pilins display a “lollipop” 3D architecture with a rounded globular head impaled on a α-helix (α1) “stick” of ∼ 50 residues [5]. The protruding N-terminal half of this stick (α1N) corresponds to a stretch of hydrophobic residues, anchoring the pilins in the cytoplasmic membrane (CM) before polymerisation. Filament assembly starts with the cleavage of the leader peptide in prepilins by the PPase, a CM-embedded aspartic protease [6]. Then, the cytoplasmic extension ATPase motor interacts with the CM-embedded platform protein [7] to extract pilins forcibly out of the membrane and polymerise them into filaments [3]. The α1-helices in pilin subunits pack roughly parallel to each other to form the filament core, exposing the globular heads on the filament surface [8–13]. In Bacteria, polymerisation is accompanied by a partial loss of α-helical order (“melting”) in α1N. The additional T4F components are involved at stages post filament assembly [14].

An important function mediated by T4F is DNA uptake, during which free DNA is bound from the extracellular milieu and translocated in vicinity of the CM [15]. This is the first step in natural transformation, a property widespread in bacteria promoting horizontal gene transfer, important for rapid evolution and the spread of antibiotic resistance [16]. In monoderm bacteria, DNA uptake is mediated by a poorly characterised T4F, the Com pilus [17]. This T4F, which has been primarily studied in *Bacillus subtilis* [18–24] and *Streptococcus pneumoniae* [25–29], exhibits a series of unique characteristics. It is a monophyletic T4F restricted to Firmicutes, almost ubiquitous in Bacilli [30]. Although its major subunit ComGC displays a typical lollipop structure, it defines a novel pilin fold, with an exclusively helical globular head [30]. Most importantly, the Com pilus appears to be much simpler than T4aP and T2SS, since it presumably consists only of core components: five pilins (ComGC, ComGD, ComGE, ComGF, ComGG), PPase (ComC), extension ATPase (ComGA) and platform (ComGB) [24].

Although the simplicity of the Com pilus could be an asset for unravelling the mechanisms of T4F assembly, our poor understanding of this filament is an obstacle. There are many important outstanding questions about Com pili. (1) Are these T4F *bona fide* pili as in *S. pneumoniae* [27, 31], or endopili as in *B. subtilis* where filaments have not been visualised on the cell surface [32]? (2) Is each of the above eight proteins required and sufficient for filament biogenesis? (3) Are ComGD, ComGE, ComGF, ComGG indeed minor pilins? (4) Do the minor pilins form a complex? In the present study, we have addressed these questions by performing a systematic analysis of the Com pilus in *Streptococcus sanguinis*. This inhabitant of the human oral cavity, which frequently causes endocarditis [33], has recently emerged as a model monoderm species for T4F [3, 17]. We finish by discuss the implications of our results for this widespread T4F superfamily of nanomachines.

## Results

### Com pili are *bona fide* T4P

Contrasting findings in *B. subtilis* [32] and *S. pneumoniae* [27, 31] make it unclear whether Com T4F are endopili or *bona fide* pili. We therefore addressed the question in *S. sanguinis*, a naturally competent species where DNA uptake is mediated by the Com pilus, like in all other Bacilli. *S. sanguinis* has recently emerged as a model monoderm species for T4F [3, 17] because it expresses T4aP, which have been extensively characterised. These T4aP, which are dispensable for competence [34], mediate twitching motility [34] and adhesion to human cells [35, 36].

Since competence in *S. sanguinis* is under the control of a peptide pheromone known as competence-stimulating peptide (CSP) [37], we imaged bacteria by transmission electron microscopy (TEM) after negative staining upon induction with synthetic CSP (the 15-aa DLRGIPNPWGWIFGR peptide). To facilitate the identification of Com filaments, we used a Δ*pil* Δ*fim* double mutant in which the two loci encoding T4aP [34] and sortase-assembled fimbriae [38] were deleted. This revealed that long surface-exposed filaments could be seen on the surface of *S. sanguinis* in the presence of CSP, with a classical T4P morphology (Fig. 1A). These filaments are ∼ 7 nm in width, ∼ 1 µm in length, and appear to be flexible. An overwhelming majority of the cells expressed a single filament.

**Fig. 1.**
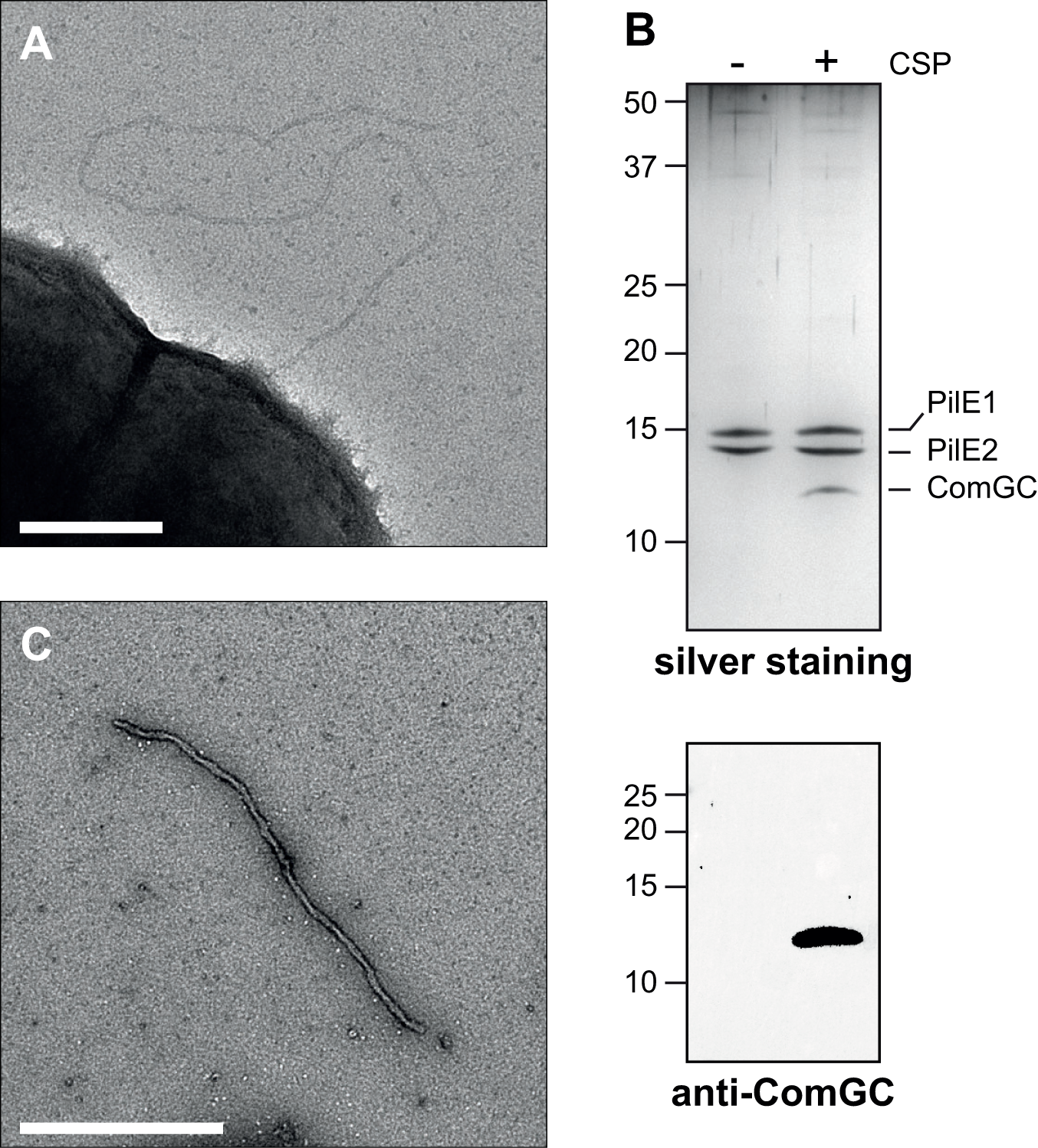
Visualisation and purification of *S. sanguinis* Com pili. **A**) Electron micrograph of negatively stained Δ*pil* Δ*fim* mutant after CSP induction. The scale bar represents 200 nm. **B**) Silver staining (upper panel) and immunoblot analysis (lower panel) of purified Com pili. Samples were prepared +/- CSP induction from cultures adjusted to the same OD_600_. Immunoblotting was performed using an anti-ComGC antibody. **C**) Electron micrograph of negatively stained purified Com pili from the *Δpil* mutant. The scale bar represents 500 nm.

Next, we designed a robust pilus purification procedure to demonstrate that the CSP-induced filaments in *S. sanguinis* are Com pili. Using an approach previously validated for *S. sanguinis* T4aP [34], we sheared filaments prepared with or without CSP by vortexing or by pipetting up and down, removed cells and debris by centrifugation, and then pelleted the filaments by ultracentrifugation. As assessed by silver staining after SDS-PAGE (Fig. 1B), although pilus preparations contained some high molecular weight contaminants, three proteins between 10 and 15 kDa were highly predominant in preparations made from the wild-type (WT) strain in the presence of CSP. In the absence of CSP (Fig. 1B), the 10 kDa protein was not seen. This smaller protein corresponds to ComGC as confirmed by immunoblotting (Fig. 1B). The two larger proteins correspond to PilE1 and PilE2, the two major subunits of *S. sanguinis* T4aP, which was expected because T4aP expression is constitutive [34]. As will be shown later, pilus preparations made from a Δ*pil* mutant consist only of ComGC.

We next used by TEM to analys pilus preparations made from the Δ*pil* mutant in the presence of CSP. An abundance of filaments could be readily seen, confirming that in *S. sanguinis* Com pili are *bona fide* pili. As shown in Fig. 1C, these filaments displayed a morphology different from those seen on cells, as they were much thicker (∼ 15 nm) and straighter. This intriguing morphological change during purification was also observed for *S. sanguinis* T4aP [34, 39]. The nature of these changes remains unexplained.

Taken together, these results show that Com pili are *bona fide* pili with a morphology typical of T4P, composed primarily of ComGC subunits.

### There is strong correlation between the transient production of the Com pilus and competence

We next determined the competence parameters in *S. sanguinis* strain 2908, where DNA transformation has not been thoroughly studied. Bacteria grown over night (O/N), were diluted 1/100 in fresh medium and incubated at 37°C. At different timepoints, from 1 to 6 h, we quantified competence by performing transformation experiments. In brief, transforming DNA (a PCR product of the *rpsL* gene in a mutant spontaneously resistant to streptomycin) and synthetic CSP were added to the bacteria. After 1 h incubation at 37°C, we counted colony-forming units (CFU) on non-selective plates and transformants on plates with streptomycin. As previously shown for other strains of *S. sanguinis* [40, 41], competence in strain 2908 was exquisitely dependent on cell density (Fig. 2A). Transformation frequency was maximal at early timepoints, reaching 8.52 ± 0.92 % at 2 h. This corresponds to 2.01 ± 0.29 10^8^ CFU/ml (Fig. 2A). Competence then decreased dramatically at higher cell densities.

**Fig. 2.**
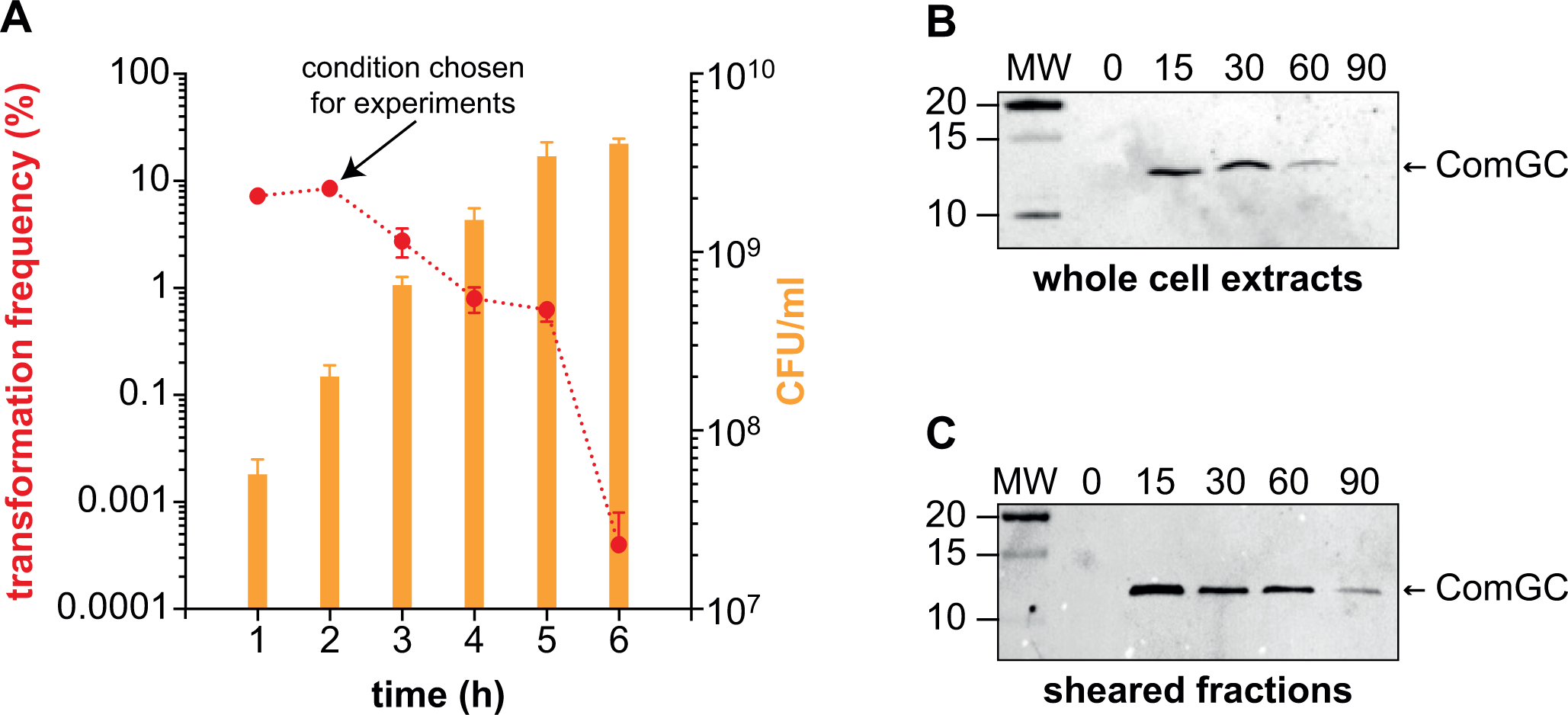
Correlation between competence, ComGC production and Com pilus production. **A**) Competence development in relation to growth phase. O/N cultures were adjusted to OD_600_ 1, diluted 100-fold in fresh THTH, and transformed after 1, 2, 3, 4, 5, or 6 h of growth at 37°C. Cell densities (CFU/ml) were determined by counting viable bacteria on non-selective plates (orange). Transformation frequencies (%) –the ratio of transformants relative to number of viable bacteria – were determined by counting transformants on plates containing streptomycin (red). The results are the average ± SD from three independent experiments. The condition chosen for further experiments – optimal transformation frequency at highest cell density – is indicated by an arrow. **B**) Immunoblot detection of ComGC in cell extracts after 0, 15, 30, 60 and 90 min induction with CSP. Extracts were quantified, equalised, and equivalent amounts of proteins were loaded in each lane. MW, molecular weight marker (in kDa). **C**) Immunoblot detection of Com T4F in sheared fractions after 0, 15, 30, 60 or 90 min induction with CSP. Fractions were prepared from equal volumes of culture, and equal volumes were loaded in each lane.

We next used immunoblotting to follow ComGC production in whole cell extracts after induction with CSP (Fig. 2B). While there was no ComGC in the absence of CSP, this protein was detected 15 min after induction. The signal peaked during the next 15 min, before then rapidly decreasing to become undetectable at 90 min (Fig. 2B). We also used immunoblotting to follow, over time, the production of surface-exposed filaments composed of ComGC (Fig. 2C). As is commonly done for T4F [42], filaments were sheared by vortexing and separated from the cells by centrifugation, the supernatants corresponding to enriched pilus fractions. After 15 min, ComGC was detected in sheared fractions, indicating that filament assembly occurs immediately after production of the major pilin subunit. As will be shown later with Δ*com* mutants, there is no cell lysis in these conditions, confirming that the protein detected in sheared fractions does not result from contamination by membrane-embedded ComGC. ComGC levels remained high in these fractions over the first 60 min, before then decreasing significantly at 90 min (Fig. 2C). Interestingly, while the time-course of Com pilus production followed that of the production of the major pilin, at 90 min ComGC was still detected in sheared fractions (Fig. 2C) while it was undetectable in whole cell extracts (Fig. 2B). These findings suggest that ComGC is stabilised upon filament assembly. Nevertheless, the Com pilus remains a short-lived T4F, which lasts only a short time.

Together, these results show that there is a strong correlation in *S. sanguinis* between the development of competence and the transient production of the Com pilus. This is comparable to what has been reported in other model species [16].

### Each *com* gene is involved in Com pilus assembly and competence

The eight *com* genes encoding this T4F in *B. subtilis* [22, 24] are present in the *S. sanguinis* genome, where they are organised in two transcription units (Fig. 3). Seven of these genes – *comGA*, *comGB*, *comGC*, *comGD*, *comGE*, *comGF*, *comGG* – form an operon [19], with overlaps between consecutive genes ranging from 1 to 41 nucleotides. The last gene *comC*, which encodes the PPase, is a distant loner [20]. This genetic organisation is conserved in the different species producing Com pili (Fig. S1). Both transcription units are preceded by a Com box (Fig. 3) [43] – TTnCGAATA – the binding site for the competence-specific σ factor ComX [44]. The expression of *comX* is under the control of (CSP) [37]. A systematic characterisation of the role of the eight *com* genes in both DNA uptake and Com pilus assembly is yet to be done. Previous studies were either focusing only on DNA uptake or were not systematic, respectively in *B. subtilis* [24] and *S. pneumoniae* [25–27]. We therefore performed such an exhaustive genetic analysis in *S. sanguinis*.

**Fig. 3.**
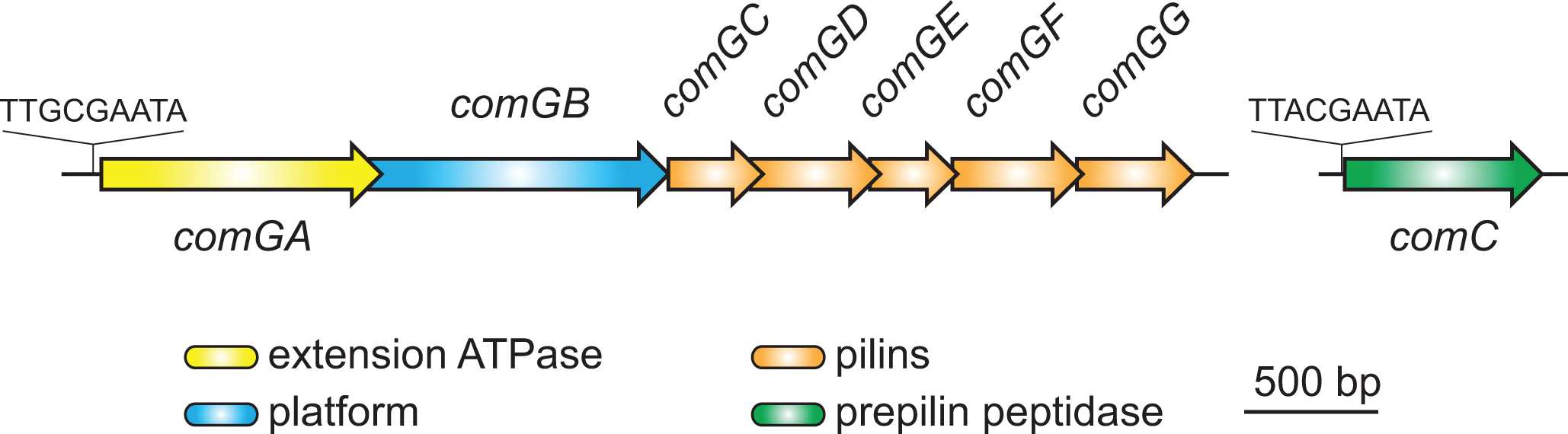
Genomic organisation in *S. sanguinis* 2908 of the eight *com* genes involved in the synthesis of the Com pilus. The Com boxes preceding both transcription units – the binding site for the competence-specific σ factor ComX – are highlighted. The genes are drawn to scale. The corresponding proteins are listed at the bottom.

Using a cloning-free mutagenesis method that generates non-polar mutants [34, 45], we deleted each of the eight *com* genes in *S. sanguinis* 2908. Target genes were replaced by a promoterless *aphA3* gene, which encodes an aminoglycoside phosphotransferase conferring resistance to kanamycin. Deletions of target genes were from start codon to ∼ 30 bp before stop codon, in order to preserve the ribosomal binding site used by the gene immediately downstream. Allelic exchange mutants were selected on kanamycin-containing plates and confirmed both by PCR and sequencing. Considering that seven *com* genes are expressed in a large operonic structure (Fig. 3), it was crucial to complement the mutants to confirm that phenotypic alterations were not due to polar effects on downstream genes. This required the design of a robust complementing platform (Fig. 4A). In brief, complementing genes were (1) put under the transcriptional control of the highly expressed lactate dehydrogenase *ldh* promoter (P*_ldh_*) [46], (2) followed by a promoterless *ermAM* gene encoding an rRNA methyltransferase that confers resistance to erythromycin, and (3) inserted in the *S. sanguinis* genome at the *pil* locus encoding T4aP [34]. The *pil* locus was entirely deleted in the process (Fig. 4A), ensuring that transformants selected on erythromycin-containing plates would be Δ*pil* mutants, to facilitate the observation of Com pili. Since the Δ*com* mutants were non-transformable (see below), construction of complemented strains was done by inserting first the P*_ldh_ com*-*ermAM* cassette *en lieu* of the *pil* locus in the WT, before deleting the corresponding *com* gene.

**Fig. 4.**
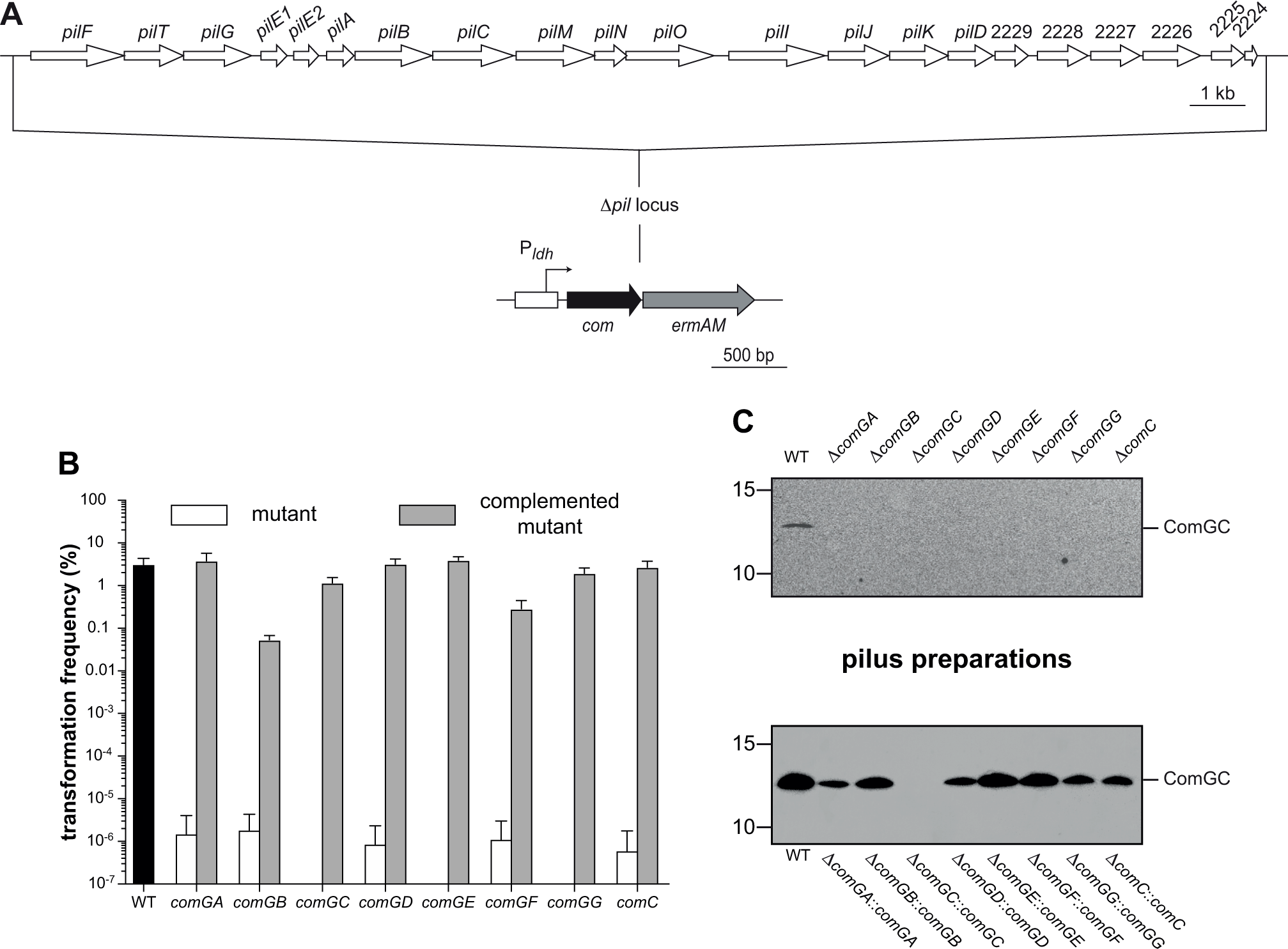
Systematic characterisation of the role of the eight *com* genes in competence and Com pilus assembly. **A**) Engineering a platform for complementation. First, we fused the complementing gene to *ermAM*, which confers resistance to erythromycin, and put them under the control of the constitutive lactate dehydrogenase promoter (P*_ldh_*) [46]. This cassette was then spliced to the upstream and downstream regions of the *pil* locus, which encodes T4aP in *S. sanguinis* [34]. The PCR product was transformed in 2908 and transformants were selected on plates containing erythromycin. As verified by sequencing, the *pil* locus was cleanly deleted and replaced by the cassette. **B**) Competence of non-polar deletion mutants in the eight *com* genes and the corresponding complemented mutants. The WT strain is included as a control. The results are the average ± SD from three independent experiments. Transformation experiments were performed as in Fig. 2. **C**) Com pilus production in non-polar deletion mutants in the eight *com* genes and in the corresponding complemented mutants. Piliation was assessed by immunoblotting using an anti-ComGC antibody on pilus preparations made from equal volumes of culture. Equal volumes were loaded in each lane. The WT strain is included as a control.

As can be seen in Fig. 4B, transformation was dramatically affected in each of the eight *com* mutants. While 3.1 ± 1.22 % of WT cells were transformed, transformation was dramatically decreased in the Δ*com* mutants (and abolished in Δ*comGC*, Δ*comGE*, Δ*comGG*), with decreases in transformation frequency of at least 6 orders of magnitude (Fig. 4B). Importantly, competence was restored, most often at WT levels, when mutants were complemented with a WT copy of the deleted gene (Fig. 4B). This demonstrates that the non-transformable phenotypes in Δ*com* mutants are not due to polar effects on downstream genes but indeed to the absence of the genes that were deleted.

We next determined whether the Δ*com* mutants were piliated by analysing pilus preparations by immunoblotting using an anti-ComGC antibody. As can be seen in Fig. 4C, pilus production was abolished in each mutant since no ComGC could be detected in pilus preparations. When mutants were complemented with a WT copy of the deleted gene, piliation was always restored except apparently for Δ*comGC*::*comGC* where no ComGC could be detected in pilus fractions (Fig. 4C). However, ComGC was produced in all strains as it could be detected in cellular extracts (Fig. S2). This important control shows that there is no contamination of pilus preparations by membrane-embedded ComGC. Critically, as seen in Fig. S2, Δ*comGC*::*comGC* is the only complemented strain in which ComGC was produced at levels much lower than in WT. This important finding shows that low-level ComGC production – too low to lead to detectable pili – allows for substantial DNA transformation.

Taken together, these results demonstrate that the eight *com* genes are required for both production of Com pili and competence for DNA transformation.

### The *com* gene are sufficient for Com pilus assembly

More than 200 genes - the *com* regulon – are under the control of CSP and ComX [43]. It was therefore formally possible that some of these other genes could be involved in the assembly of Com pili. We therefore endeavoured to engineer a strain that would express the eight *com* genes constitutively to determine whether that could lead to the production of Com pili. We constructed such a strain in two steps (Fig. 5A). First, we used allelic exchange to swap the promoter in front of the seven-genes *com* operon with P*_ldh_* (generating the intermediate strain P*_ldh_ comG*). Then, we inserted in this intermediate strain a P*_ldh_ comC*-*ermAM* cassette at the *pil* locus (generating the final strain P*_ldh_ comG* P*_ldh_ comC*) (Fig. 5A).

**Fig. 5.**
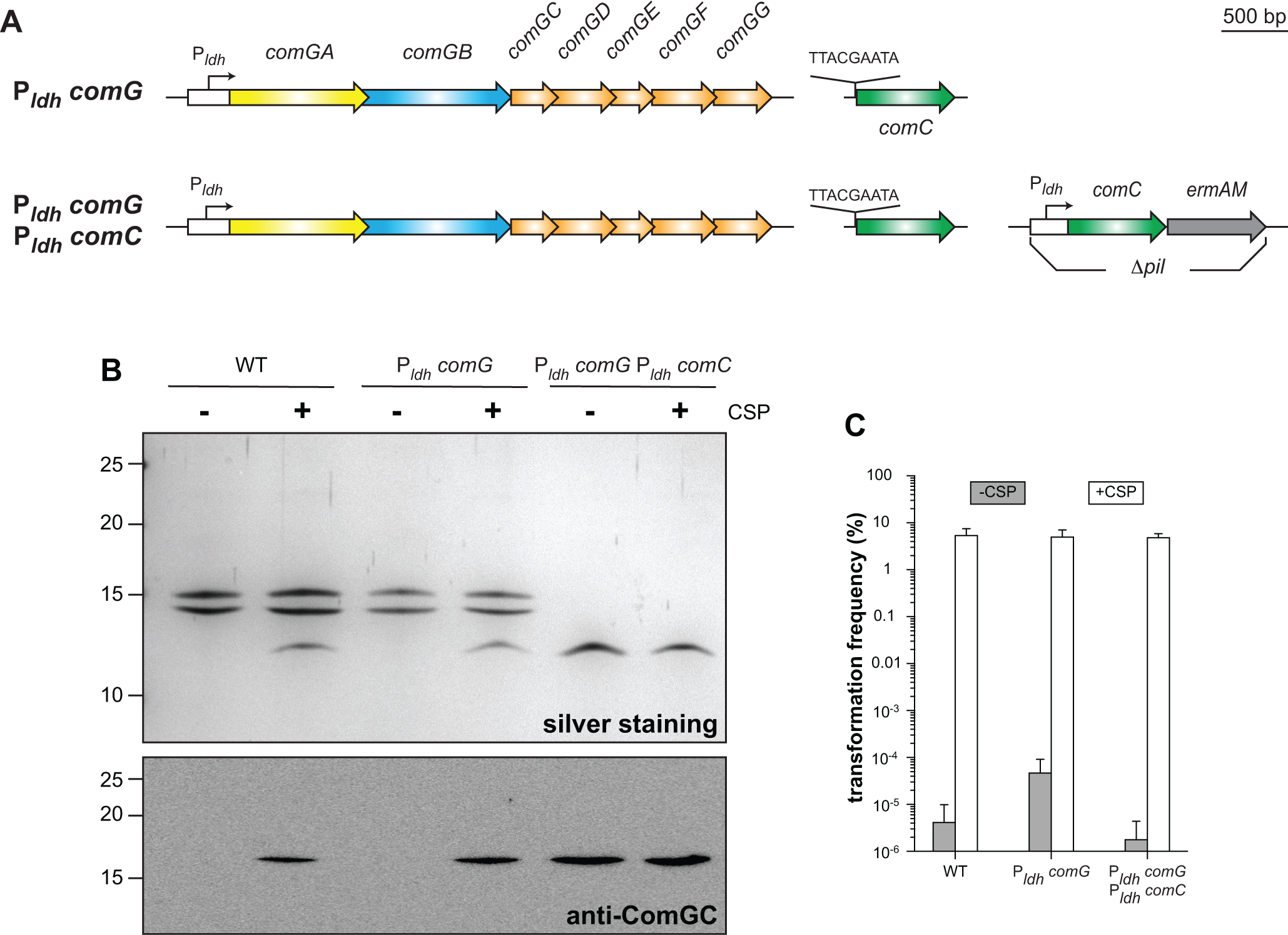
The eight *com* genes are sufficient for Com pilus assembly. **A**) Engineering a strain expressing Com pili constitutively. First, we used allelic exchange to replace the promoter of the seven-genes *com* operon by P*_ldh_*, generating the intermediate strain. We then inserted a P*_ldh_ comC*-*ermAM* cassette *en lieu* of the *pil* locus, generating the final strain. **B**) Silver staining (upper panel) and immunoblot analysis (lower panel) of purified Com pili from WT, intermediate/final strains. Samples were prepared +/- CSP induction from cultures adjusted to the same OD_600_. Immunoblotting was performed using an anti-ComGC antibody. Molecular weight markers (in kDa) are indicated on the left. **C**) Competence of WT and intermediate/final strains ± CSP. The results are the average ± SD from three independent experiments. Transformation experiments were performed as in Fig. 2.

We showed by immunoblotting on cellular extracts that ComGC production became independent of CSP in the intermediate strain (Fig. S3). However, the pilin was not processed in the absence of CSP until *comC* expression became independent of CSP as well (Fig. S3). Pilus preparations, analysed by silver staining and immunoblotting, showed that the P*_ldh_ comG* P*_ldh_ comC* expressed Com pili constitutively (Fig. 5B). No T4aP were present in the final strain because of the deletion of the *pil* locus. In contrast, in the intermediate strain P*_ldh_ comG*, Com pili were produced only in the presence of CSP (Fig. 5B) because the pilin was not processed in the absence of CSP (Fig. S3). Finally, we tested competence by measuring transformation frequencies (Fig. 5C). We found that the Com pili produced by the final strain – although they are produced constitutively – are capable of mediating transformation to WT-levels (Fig. 5C), showing that they are perfectly functional. However, since the *com* regulon is > 200 genes [43], CSP was still needed for competence.

Together, these results show that the eight *com* genes are not only necessary but also sufficient for Com pilus assembly.

### The Com pilus is a minimalistic T4F consisting only of core components

It was previously noted that the Com proteins in *S. sanguinis* display sequence conservation with orthologs in the model species *B. subtilis* and *S. pneumoniae* [30]. Here, we performed a detailed bioinformatic analysis to determine whether Com proteins correspond to one of the four core components universally conserved in T4F [3].

We identified known protein domains using primarily InterPro [47] and predicted 3D structures using AlphaFold [48] (Fig. 6). All AlphaFold-generated models exhibited high confidence scores (Fig. S4). In brief, ComGA is the extension ATPase, the cytoplasmic motor undergoing conformational changes upon ATP hydrolysis to power T4F assembly [7]. ComGA adopts a typical hexameric ring-shaped 3D structure, with a central pore (Fig. 6A). ComGB is the platform, a membrane-embedded T4F assembly hub [3] transmitting mechanical forces generated by the extension ATPase to membrane-embedded pilins. ComGB is predicted to exhibit a “cherry” pair-like structure [3], where each cherry corresponds to a repeated globular domain (Fig. 6B). ComC is the PPase, an aspartic protease that processes prepilins [6]. ComC has a bi-modular structure with a C-terminal domain involved in the proteolysis of the leader peptide in prepilins, and an N-terminal domain involved in N-methylation of mature pilins (Fig. 6C) [49]. The last five proteins – ComGC, ComGD, ComGE, ComGF, ComGG – are type 4 pilins, all displaying N-characteristic terminal class 3 sequence peptides (SP3) (Fig. S5) [5, 50]. The hydrophilic leader peptides – cleaved by the PPase – are 8-15 residues in length and invariably end with an Ala. ComGC is the major pilin [32], while the remaining four proteins are likely to be minor pilins, which is strengthened by AlphaFold predictions that these proteins all display characteristic lollipop structures (Fig. 6D). It should be noted that the extreme C-terminus of ComGG is the only portion with a very low confidence score (Fig. S4). Interestingly, none of these minor pilins has a purely helical globular head, which thus remains a distinctive feature of ComGC [30].

**Fig 6.**
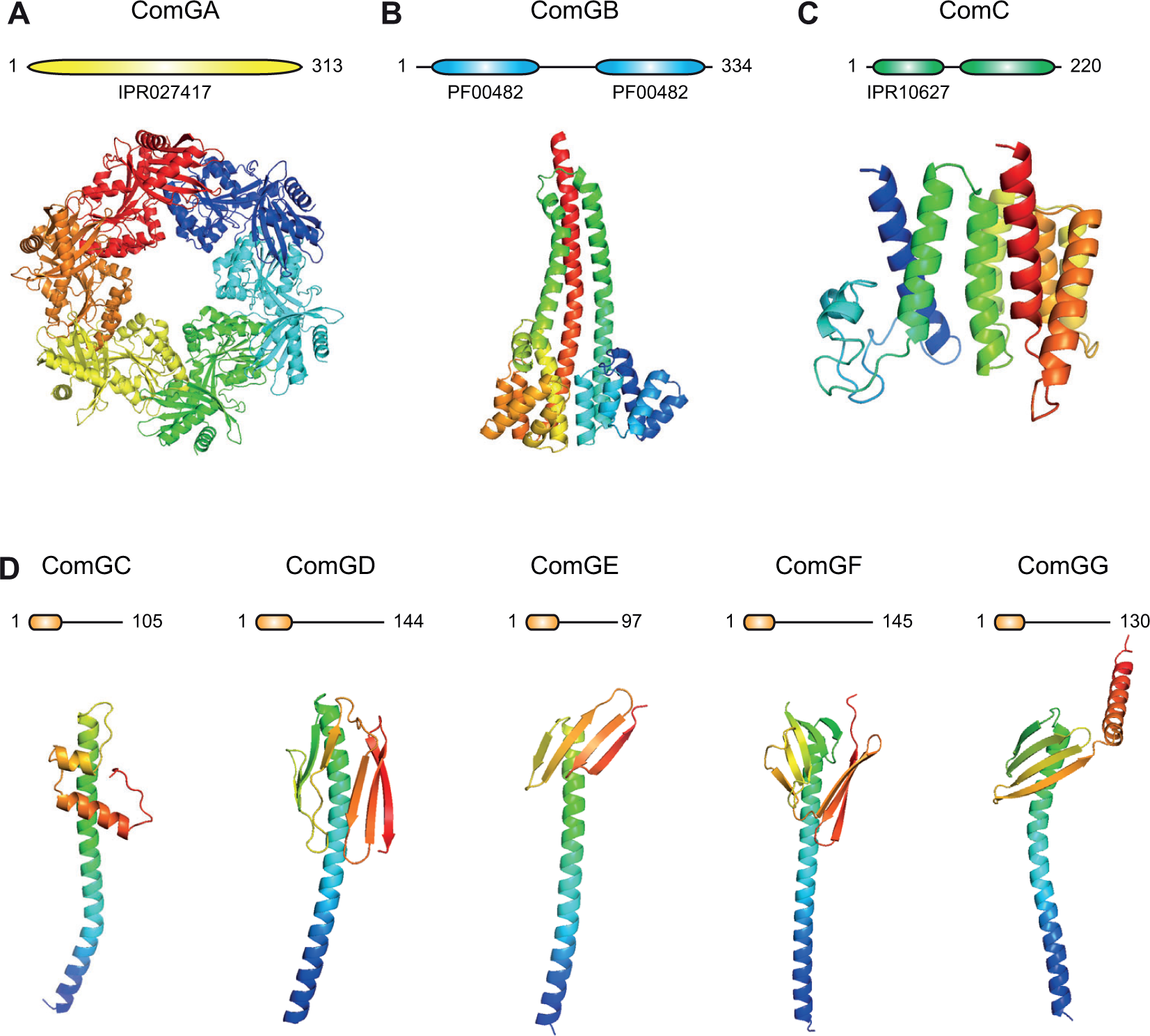
The Com machinery in *S. sanguinis* is a minimalistic T4F composed only of core proteins. Bioinformatic analysis of each protein: protein architecture drawn to scale (upper panel) and AlphaFold predicted structure (lower panel) in cartoon view, rainbow-colored from blue (N-terminus) to red (C-terminus). **A**) ComGA is the extension ATPase with an IPR027417 domain. The predicted structure is for the typical hexameric ring with a central pore **B**) ComGB is the platform protein with two PF00482 domains. **C**) ComC is the PPase with two domains: an N-terminal methylase (IPR10627) and a C-terminal peptidase. **D**) ComGC, ComGD, ComGE, ComGF and ComGG are type 4 pilins, represented under their processed form. They all start with a protruding α1N (orange rounded boxes) and display lollipop 3D architectures.

Taken together, these findings unambiguously show that the Com machinery is composed only of core components universally conserved in T4F. The Com pilus is thus a minimalistic T4F, significantly simpler than traditional T4aP and T2SS models.

### ComGD, ComGE, ComGF and ComGG are minor pilins present in Com pili in very low abundance

Next, we focused on the five pilins in the Com system, which display an N-terminal SP3 (Fig. S5). The major pilin ComGC served as a positive control. We first tested whether ComGD, ComGE, ComGF and ComGG are indeed *bona fide* minor pilins, *i.e.*, processed by the prepilin peptidase ComC and incorporated in the pili as low abundance subunits. After generating rabbit antisera against these five proteins, we detected them by immunoblotting in the WT strain and a Δ*comC* mutant in which the PPase production is abolished. In the WT strain, processing by ComC, which removes the leader peptide in pilins, is expected to generate mature proteins shorter than in the Δ*comC* mutant. As seen in Fig. 7, in *ΔcomC*, we detected proteins slightly longer than in the WT corresponding to unprocessed precursors, confirming that ComGD, ComGE, ComGF and ComGG (as well as ComGC) are all cleaved by the PPase ComC. The detection of a methylase domain in ComC (Fig. 6C) suggests that cleavage of the leader peptide will be accompanied by methylation of the new N-terminus, as in many other bacterial T4F [49].

**Fig. 7.**
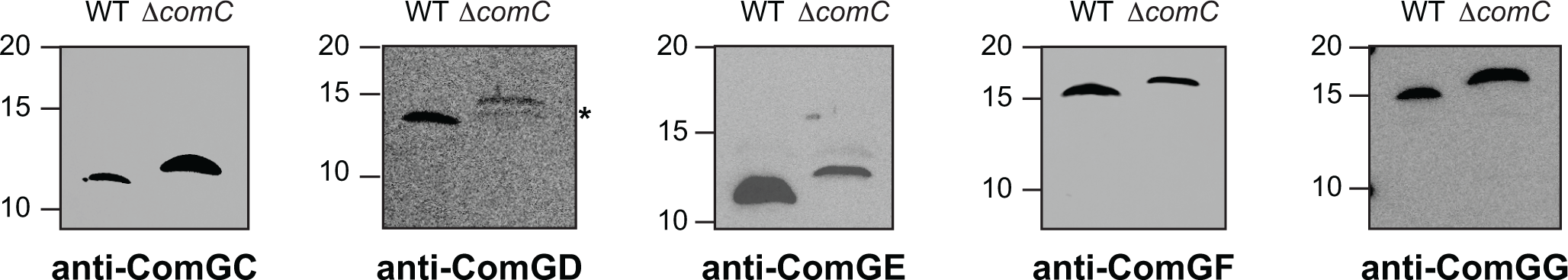
ComGC, ComGD, ComGE, ComGF and ComGG are genuine type 4 pilins processed by the PPase ComC. Immunoblot analysis of processing by ComC of the leader peptide in the five Com pilins. We used specific antibodies, which were generated for this study. Whole cell protein extracts were quantified, equalised, and equivalent amounts of proteins were loaded in each lane. *Spurious band (or degradation product) detected using the anti-ComGD antibody, which was of limited quality. Molecular weight markers (in kDa) are indicated on the left.

Unfortunately, our attempts to detect ComGD, ComGE, ComGF and ComGG in pilus preparations by immunoblotting were repeatedly unsuccessful, which is not unexpected. Indeed, WT pilus preparations are poorly concentrated because Com pili are produced at very low bacterial density, one filament/cell, and only during a short time window [16]. Therefore, since they are predicted to be present only at one copy/filament, the minor pilins are too scarce in these preparations and go undetected in contrast to the major pilin ComGC. The pilus preparations in the final strain P*_ldh_ comG* P*_ldh_ comC* were more concentrated than those made from the WT (Fig. 5B) because they could be produced at much higher cell density. We therefore used these pilus preparations to test again by immunoblotting whether ComGD, ComGE, ComGF and ComGG could be detected. As can be seen in Fig. 8, we could detect all four pilins in concentrated pilus preparations, showing that they are indeed low abundance components of the Com pili and thus genuine minor pilins. As a negative control, we showed that these minor pilins were not detected in pilus preparations made from the intermediate strain P*_ldh_ comG*, which did not produce Com pili in the absence of CSP (Fig. 5B).

**Fig. 8.**
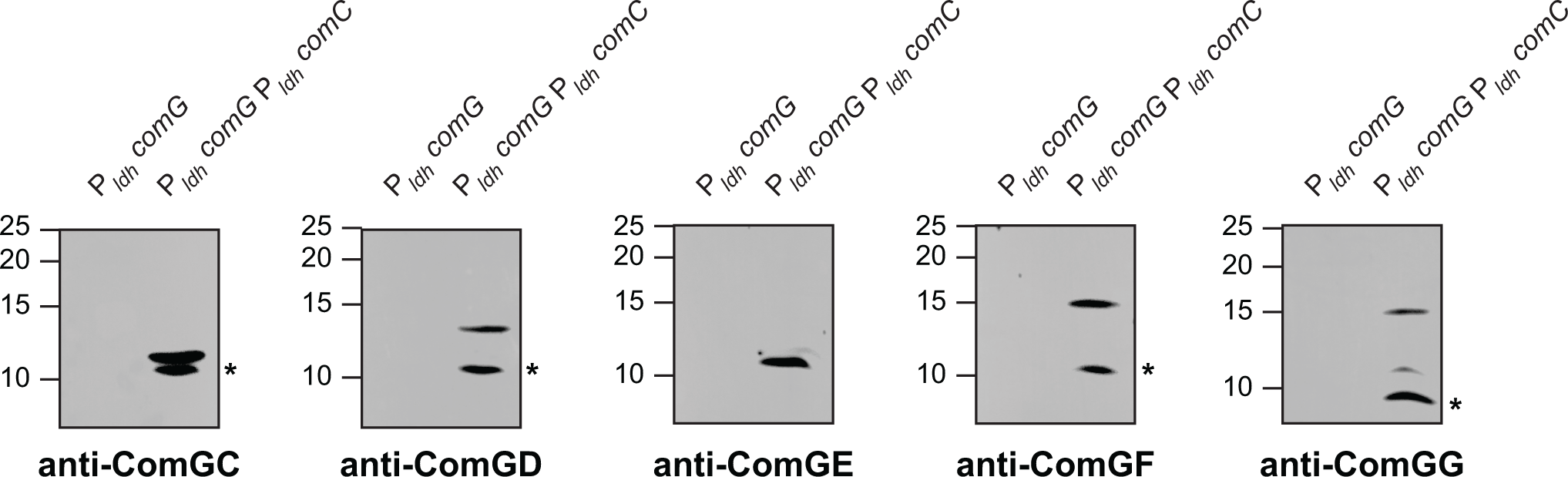
ComGD, ComGE, ComGF and ComGG are minor pilus components. Immunoblot analysis of the co-purification of the five pilins (ComGC is a positive control). Pilus preparations from the final strain (expressing Com pili constitutively) and the intermediate strain (non-piliated) were analysed. *Spurious band at 10 kDa detected with several antibodies, corresponding to an unknown protein. Molecular weight markers (in kDa) are indicated on the left.

Taken together, these results show that Com pili are composed of five pilin subunits. While ComGC is the major subunit, ComGD, ComGE, ComGF and ComGG are four minor pilins, present in the filaments in very low abundance.

### The four minor pilins form a complex predicted to be at the pilus tip

T4aP and T2SS both contain a set of four minor pilins – often named with the letters H, I, J, K – essential for filament assembly [51, 52]. These minor pilins interact to form a tip-located complex [53]. Since T4F are assembled from tip to base, filament assembly must start with this complex of minor pilins [52].

We therefore characterised the interactions between the four minor Com pilins. Since interacting proteins often exert a stabilising effect on each other [45], we used immunoblotting to assess the stability of every pilin in non-polar deletion mutants (Fig. 9). Critically, each protein (1) was detected in the WT strain, (2) undetectable in a non-polar mutant in which the corresponding gene was deleted, and (3) its production was restored upon complementation. As seen in Fig. 9, several pilins displayed reduced levels in the absence of others, which suggests that they interact. We ruled out the possibility that the above effects could be due to polar effects since complementation with a WT copy of the mutated gene restored the stability of the interacting partners (Fig. 9). In summary, ComGC was mainly unaffected by the absence of other pilins and had no major impact on their stability. ComGD stabilised ComGE and was stabilised by ComGE and ComGF. ComGE and ComGF showed mutually stabilising effects – they were strongly dependent on each other for stability – and the absence of either protein resulted in reduced levels of ComGG. Finally, ComGG had no detectable impact on the stability of other pilins.

**Fig. 9.**
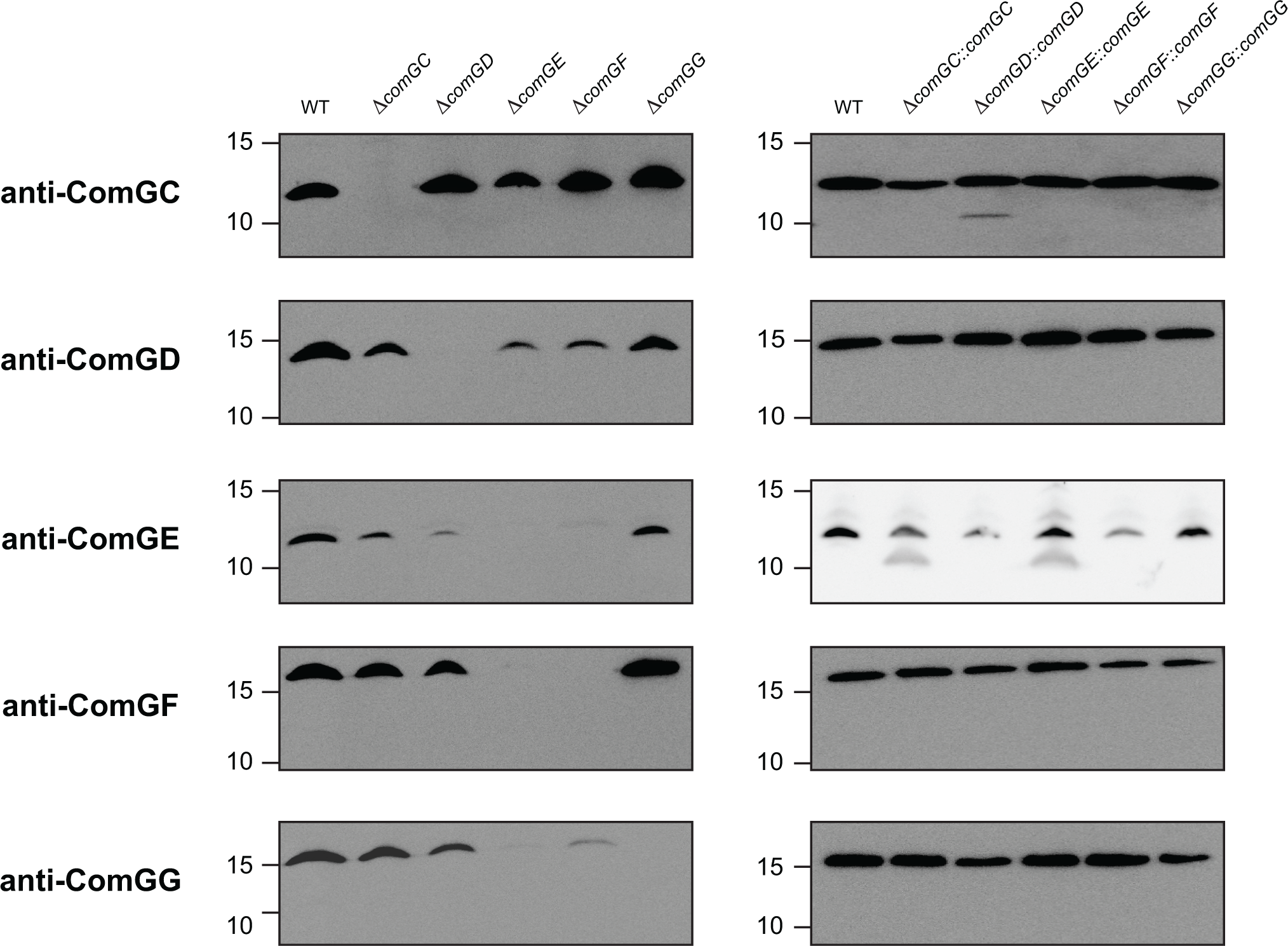
The four minor pilins interact to form a complex. Immunoblot analysis of the stability of each pilin in whole-cell protein extracts of pilin mutants and mutants complemented with a WT copy of the deleted gene. The WT strain was included as a positive control. Extracts were quantified and equivalent amounts of proteins were loaded in each lane. Molecular weight markers (in kDa) are indicated on the left.

In other T4F, the four interacting minor pilin subunits form a tip-located H-J-I-K complex [53], capped by the K subunit lacking the conserved Glu_5_ [54, 55] (Fig. S6A). Since ComGD, ComGE, ComGF and ComGG show structural and sequence similarities to the H, I, J, K subunits, we modelled the D-F-E-G complex using AlphaFold (Fig. 10). This complex, which has a high confidence score (Fig. S4), is capped by ComGG, which is the only subunit lacking the conserved Glu_5_. The next subunit is ComGE, followed by ComGF, which likely explains their mutually stabilising effects. The bottom subunit is ComGD, which is predicted to be the adapter between the F-E-G complex, and the filament formed by ComGC. The D-F-E-G model is consistent with the interaction data (Fig. 9), and structurally similar to the H-J-I-K complex in T2SS [55] (Fig. S6B).

**Fig. 10.**
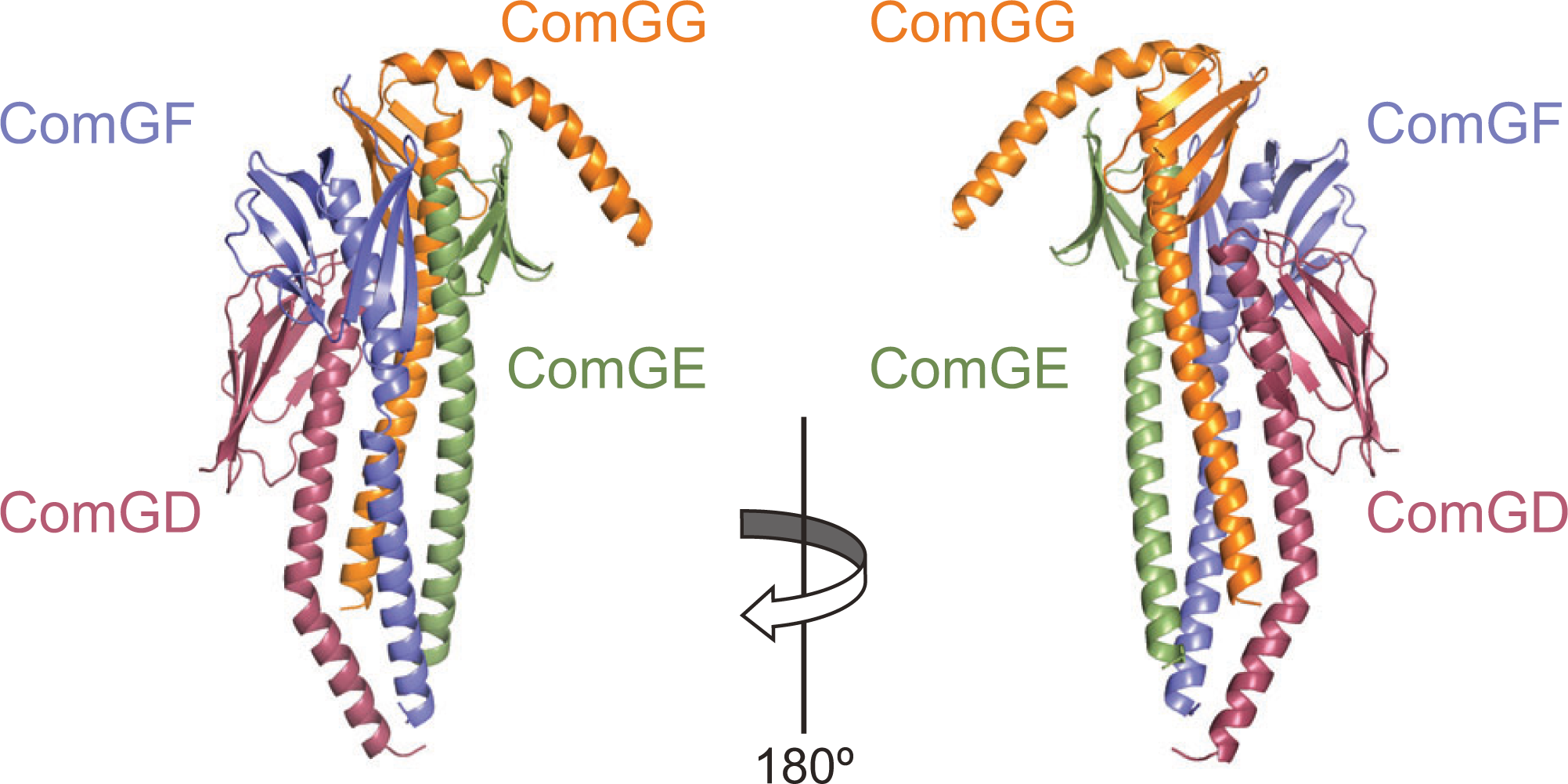
Structural model of the ComGD-ComGF-ComGE-ComGG complex of four minor pilins, predicted to be located at the tip of Com pili. The AlphaFold-predicted structure is presented in cartoon 180° views, with the four subunits highlighted in different colours.

Taken together, these results show that Com minor pilins interact, exerting strongly stabilising effects on one another. The resulting complex could be modelled in the order ComGD-ComGF-ComGE-ComGG and is predicted to be at the tip of Com pili.

## Discussion

T4F are a superfamily of exceptionally versatile nanomachines, ubiquitous in Bacteria and Archaea, that use conserved multi-protein machineries to assemble and operate filaments composed of type 4 pilins [1, 2]. Although T4F have been studied for more than 40 years – because they mediate important biological functions and are key virulence factors in numerous bacterial pathogens – the mechanisms of filament assembly and/or T4F-mediated functions remain poorly understood. One of the main underlying reasons is that the best characterised T4F – T4aP and T2SS – are very complex nanomachines, with the largest number of protein parts. Therefore, in this report we took advantage of the inherent simplicity of T4F in monoderm bacteria [17] to systematically characterise one of the arguably simplest T4F: the Com pilus that mediates DNA uptake in naturally competent monoderm bacteria. This leading to the findings discussed below, shedding light on important aspects of T4F biology.

Using *S. sanguinis*, which has emerged as a monoderm T4F model in the last decade [3, 17], we show that Com filaments are *bona fide* surface-exposed pili with a morphology typical of T4P, rather than elusive endopili. This finding is important for several reasons. First, it is coherent with the notion that T4F act as molecular harpoons capturing free DNA from the environment to initiate DNA uptake [56]. Second, there was uncertainty about Com pili as they have been described as pili in *S. pneumoniae* [27, 31] and endopili in *B. subtilis* [32]. Third, pili and endopili although they are extremely different types of filaments can be assembled by almost identical T4F machineries such as T4aP and T2SS. Fourth, our findings confirm that Com pili, which are a phylogenetic clade of T4F with specific genes profile and organisation [1], are genuine T4P corresponding to a new subtype. As suggested recently [3], Com pili should be classified as T4dP from now on. Critically, the finding that competence was restored in the complemented mutant Δ*comGC*::P*_ldh_ comGC* at pilin levels too low for pili to be detected or visualised – which mirrors findings with *N. gonorrhoeae* T4aP [57], which are involved in DNA uptake in this diderm species – offers an explanation for the paradox that surface-exposed Com pili have not yet been seen in *B. subtilis*. Indeed, it is possible that ComGC levels in the 168-derived model *B. subtilis* strain are too low for the assembly of surface-exposed µm-long pili, but enough for the assembly of very short (and thus undetectable) filaments allowing substantial DNA transformation. It would be interesting to test whether surface-exposed Com pili could be detected in other *B. subtilis* isolates, or in derivatives of strain 168 in which a stronger promoter would be used to drive the expression of the *com* genes.

The finding that Com pili are a minimal T4F only composed of core components has crucial implications for the understanding of the biology of these filaments. Our results provide strong experimental evidence for the hypothesis that pilin, PPase, extension ATPase and platform protein – the only four components found in every T4F system – constitute the minimal machinery necessary for filament assembly [3]. Indeed, we show that the monoderm *S. sanguinis* uses only these four core components to produce a surface-exposed µm-long pilus dedicated to DNA uptake, the Com pilus. Until a system with fewer minor pilins (or even none) is discovered – if such a system exists – the T4dP remain the simplest known T4F, possibly the closest to what the original T4F might have looked like [1]. It lends support to the following, generally applicable, mechanistic model of T4F assembly. The prepilins are translocated across the CM by the Sec machinery [58, 59], adopting the topology necessary for their processing by the PPase. Prepilins are anchored in the CM via the stretch of hydrophobic residues in their protruding α1N stick, with their leader peptide in the cytoplasm, and their globular heads outside [60]. This applies to each of the five Com prepilins, which all have SP3 and are structurally predicted to be lollipops. The leader peptide in prepilins is then cleaved by the PPase ComC, a CM-embedded aspartic protease. Interestingly, the finding that ComGC is not processed in the intermediate strain P*_ldh_ comG* expressing the *comG* operon constitutively (but not *comC*), indicates that the second PPase PilD involved in biogenesis of T4aP cannot recognise Com pilins. As proposed recently [3], processed pilins are assembled into filaments by a spindle motor composed of the extension ATPase and platform. The platform, which acts as a “drill chuck” holding a pilin, is powered by the rotation of the hexameric extension ATPase. It loads pilin subunits, extracts them from the membrane – which is accompanied by a partial melting of α1N – and polymerises them in a helical array with the α1-helices at the core.

The last notable finding in this study is that despite its simplicity, the Com nanomachine comprises four broadly conserved minor pilins required for piliation, forming a complex at the tip of the pilus. The ComGD-ComGF-ComGE-ComGG complex is homologous to similar complexes in T4aP (PilH-PilJ-PilI-PilK) and T2SS (GspH-GspJ-GspI-GspK) [54, 55]. These four pilins sets shared several features (1) they are encoded by an operon in the order HIJK (DEFG in the Com system), (2) the K subunit (ComGG in the Com system) is the only one lacking a Glu_5_ residue, (3) they form a complex in the order HJIK (DFEG in the Com system), predicted to be located at the tip of the filaments (this was recently confirmed by cryo-electron tomography in T4aP [53]). The complex is capped by the K subunit (ComGG in the Com system), which is coherent with the lack of Glu_5_, a residue usually allowing neutralisation of the positively charged N-terminus of the prior subunit in the pilus (counted from tip). The broad conservation of this tip-located complex of four minor pilins in very different T4F is in favour of a key role. Although this role is yet to be experimentally determined, it is likely to be broadly conserved since it has been shown that the HIJK set from T2SS can promote T4aP assembly [52]. However, these minor pilins might also play specific roles in different systems as suggested by (1) significant structural differences in the filament-capping subunits (GspK has a large extra module grafted on its globular head, while the C-terminus in ComGG is a region without confident prediction), and (2) the finding that PilHIJK interact with the PilC/PilY1 adhesin [53] and GspHIJK are involved in the recognition of secreted effector [61]. Perhaps, DEFG could be involved in DNA binding by Com pili.

In conclusion, by performing a systematic analysis in *S. sanguinis* of a bacterial T4F involved in DNA uptake in hundreds of naturally competent monoderm species, we have shown that Com pili – representing a novel T4dP subtype – correspond to a minimalistic T4F, with less protein parts than previously studied ones. Our findings have not only important implications for all T4F – most notably about the mechanisms of filament assembly – but they pave the way for further investigations in a minimalistic T4F model with dramatic potential to improve our understanding of a mesmerising superfamily of nanomachines playing a key role in prokaryotic biology.

## Material and methods

### Bacterial strains and growth conditions

*E. coli* DH5α was used for cloning. It was grown in liquid or solid lysogeny broth (LB) medium (Difco), containing, when required, 100 µg/ml spectinomycin (Sigma). Chemically competent cells were prepared as described [62]. Overall, we used standard molecular biology techniques [63]. If needed, PCR products were cloned directly into pCR8/GW/TOPO (Invitrogen), as instructed by the manufacturer, and/or purified using a QIAquick PCR purification kit (Qiagen).

Todd Hewitt (TH) broth (Difco) was used to grow *S. sanguinis* as described [34], with minor modifications. TH plates, made upon addition of 1.5 % agar (Difco), were incubated at 37°C in anaerobic jars (Oxoid) under anaerobic conditions generated using Anaerogen sachets (Oxoid). Liquid cultures – containing when required 0.05 % tween 80 (Merck) (this is called THT) to limit bacterial clumping, and/or 100 mM HEPES (Euromedex) (this is called THTH) to prevent acidification of the medium – were grown statically under aerobic conditions. When required, kanamycin (500 μg/ml), erythromycin (5 μg/ml), or streptomycin (100 μg/ml) was used for selection. When needed, 15 mM *p*-Cl-Phe (Sigma), was used for counterselection.

All the *S. sanguinis* strains that were used in this study (Table S1) are derivatives of the fully sequenced 2908 throat isolate [34]. Strains were constructed as follows. Genomic DNA was prepared from liquid cultures using the XIT Genomic DNA from Gram-Positive Bacteria kit (G-Biosciences), as instructed by the manufacturer. PCR were done using high-fidelity DNA Polymerase (Agilent). Marked non-polar deletion mutants were constructed using a splicing PCR (sPCR) strategy involving no cloning [34, 45]. For each target gene, three PCR products were generated with *aphA3*-F/*aphA3*-R, Δ-F1/Δ-R1 and Δ-F2/Δ-R2 pairs of primers (Table S2), spliced by PCR and transformed directly into *S. sanguinis* (see below). During this process, target genes were deleted and replaced by a promoterless *aphA3* gene conferring resistance to kanamycin. Genomic DNA extracted from Km^R^ transformants was tested by PCR and sequencing, to confirm that the desired allelic exchange events had taken place. The unmarked Δ*pil* deletion mutant was constructed by sPCR using a previously described two-step methodology [64], which uses a mutant *pheS* as a counterselectable marker. Primary Km^R^ mutants in which the targe gene was deleted and replaced by a *pheS* aphA-3* double cassette were transformed with a sPCR product in which the regions upstream and downstream the target gene were fused by PCR and the Km^S^ unmarked Δ*pil* mutants were counterselected on plates containing 15 mM *p*-Cl-Phe. The unmarked Δ*fim* mutant was constructed in one step by transforming *S. sanguinis* directly with a sPCR product in which the regions upstream an downstream the target gene were fused. We used a protocol optimised for identification of transformants in the absence of selective pressure (see below). Transformants were identified by PCR and sequencing.

Mutants were complemented by designing a new complementation platform, originally with the *comX* gene. Using sPCR (Table S2), we put the promoterless *comX* under the transcriptional control of the *S. sanguinis* promoter of lactate dehydrogenase (P*_ldh_*), which is highly and constitutively expressed in streptococci [46]. This was cloned into pCR8/GW/TOPO. The P*_ldh_ comX* cassette was then fused, by sPCR, to a promoterless *ermAM* gene conferring resistance to erythromycin. This was cloned into pCR8/GW/TOPO. Finally, the regions flanking the *pil* locus encoding T4aP [34] were spliced to this P*_ldh_ comX-ermAM* cassette, and transformed directly into *S. sanguinis*, replacing the *pil* locus in the process. Genomic DNA extracted from Ery^R^ transformants was tested by PCR and sequencing, to confirm that the desired allelic exchange events had taken place. Subsequently, this construct was used to generate three PCR products for each complementing gene with compF/compR, compF1/compR1 and compF2/compR2 pairs of primers (Table S2), which were spliced by PCR and transformed directly into *S. sanguinis* (see below). Ery^R^ transformants were verified by PCR and sequencing.

To engineer a strain expressing Com pili constitutively, we first generated the P*_ldh_ comG* intermediate strain in which the endogenous promoter of the *comG* operon was replaced by P*_ldh_*. We amplified the P*_ldh_ comGA* locus from the corresponding complemented strain, to which we spliced to the region upstream the endogenous promoter of the *comG* operon. This sPCR product was transformed directly into *S. sanguinis* using a protocol optimised for identification of transformants in the absence of selective pressure. Transformants were identified by PCR and sequencing. Then, we transformed this intermediate P*_ldh_ comG* strain with the P*_ldh_ comC*-*ermAM* cassette from the corresponding complemented strain. Ery^R^ transformants, verified by PCR and sequencing, corresponded to P*_ldh_ comG* P*_ldh_ comC* in which all the genes necessary for the biogenesis of the Com pilus were under the control of the constitutive P*_ldh_* promoter. The final strain thus has two copies of *comC*.

### Transformation assays

*S. sanguinis* was routinely transformed as described [34], with minor modifications. In brief, bacteria grown O/N in THTH were diluted to OD_600_ 0.01 in pre-warmed THTH and incubated at 37°C for 2 h. We then induced competence in this culture by adding synthetic CSP at a final concentration of 227 ng/ml, took a 330 µl aliquote to which we added 300 ng transforming DNA corresponding. After incubation for 1 h at 37°C, bacteria were bath-sonicated for 1 min at medium amplitude on a Bioruptor (Diagenode) to break bacterial chains and plated on plates with suitable antibiotics to select for transformants.

For quantifying competence, a similar protocol was used. After O/N growth, bacteria were diluted in THTH. Induction was performed with 227 ng/ml CSP, and 100 ng purified *rpsL* PCR product from a 2908 mutant spontaneously resistant to streptomycin was used [34]. After bath-sonicating for 45 s, we performed serial dilutions that were plated and grown on non-selective plates for counting CFU, and on plates with streptomycin for counting transformants. Transformation frequencies (in %) were determined as total number of transformants/total CFU. We used a very similar protocol for non-selective transformation, the only major difference being that bacteria were diluted 10^-7^ in THTH after O/N growth. If analysed colonies obtained on plates with no antibiotics were non-clonal, as verified by sequencing, the bath-sonication was repeated the next day.

### Visualisation and purification of Com pili

Surface-associated Com pili in *S. sanguinis* were visualised by transmission electron microscopy after negative staining, as follows. After O/N growth, bacteria were diluted in THTH to OD_600_ 0.01 and grown to OD_600_ 0.04-0.08, before pilus production was induced 20 min with 300 mg/ml CSP. Bacteria were adsorbed for 3 min to glow-discharged carbon-coated grids (EMS) and fixed 5 min in 2 % glutaraldehyde. The grids were floated on a drop of pilus buffer (20 mM Tris, pH 7.5, 50 mM NaCl), which was repeated 10 times, and then stained for 2 min with 2 % aqueous uranyl acetate. Stain solution was gently drained off the grids, which were air-dried before visualisation using a Tecnai 200KV electron microscope (Thermo Fisher Scientific). Digital image acquisition was made with a numeric camera (16 megapixel, CMOS, Oneview, Gatan).

Com pili were purified from *S. sanguinis* by adapting a protocol previously used to purify *S. sanguinis* T4aP [34]. Liquid cultures grown O/N in THTH were used to re-inoculate pre-warmed THTH at OD_600_ 0.01 and grown statically for 2 h. CSP was then added at a final concentration of 300 ng/ml and induction was performed during 30 min. Bacteria were pelleted by centrifugation for 15 min at 4,149 *g* at 4°C. Pili were sheared after re-suspending bacterial pellets in ice-cold pilus buffer either by vortexing 1 min at full speed, or by repeated pipetting up and down. Bacteria were then pelleted by two rounds of centrifugation at 4°C for 10 min at 9,220 *g*.

Occasionally, the supernatant containing the pili was filtered using 0.2 μm syringe filters (Sigma-Aldrich). Finally, pili were pelleted by ultracentrifugation at 100,000 *g* for 1 h at 4°C. The pellets were resuspended in pilus buffer by pipetting up and down.

### Preparation of protein extracts, SDS-PAGE, antisera, and staining/immunoblotting

We prepared whole cell protein extracts as follows. Liquid cultures grown O/N in THTH were used to re-inoculate pre-warmed THTH at OD_600_ 0.01 and grown statically for 2 h. When specified, CSP was then added at a final concentration of 300 ng/ml and induction was performed during 30 min. Bacteria were pelleted by centrifugation at 4°C for 15 min at 4,149 *g*, or 20 min at 9,220 *g*. Pellets were re-suspended in 1 ml ice-cold pilus buffer, transferred to Lysing Matrix B tubes (MPBio) and lysed in a FastPrep-24 5G (MPBio) using five cycles (1 min on at 6,5 m/s, 1 min off). In some experiments, pili were sheared and removed by centrifugation as described above, before the protein extracts were prepared (we refer to these as cellular extracts).

SDS-PAGE was carried out in a Mini-Protean Tetra cell system (Bio-Rad) for 1 h at 200 V, using either Tris-Glycine (Bio-Rad) or Tris-Tricine buffers [65]. The Precision Plus Protein All Blue Prestained Protein Standards (Bio-Rad) was used as molecular weight marker. Gels were either stained – with Bio-Safe Coomassie (Bio-Rad) or Pierce Silver Stain Kit (Thermo Scientific) – or transferred to a membrane and analysed by immunoblotting.

Polyclonal antisera against ComG pilins were produced by immunising rabbits with a mixture of two different peptides that were synthesised from each protein (Eurogentec). Peptides corresponding to the following residues in mature pilins were used: ComGC (64-78 and 80-94), ComGD (87-101 and 115-129), ComGE (60-74 and 72-86), ComGF (80-94 and 122-136), and ComGG (62-76 and 91-105). Immunoblotting was done as follows. After SDS-PAGE, proteins were transferred onto Amersham Hybond ECL nitrocellulose membrane (GE Healthcare). The wet transfer was carried out for 1 h at 100 V in ice-cold buffer (39 mM glycine, 48 mM Tris base, 0.037 % SDS, 20 % isopropanol). The blotted membranes were blocked for 1 h at room temperature, while shaking, in PBS supplemented with 0.1 % Tween-20 (PBST) containing 5 % (w/v) skim milk powder (VWR). The membranes were then incubated for 1 h with primary antibodies diluted in PBST with or without 5 % milk: 1/2,500 (anti-ComGC) or 1/1,000 (anti-ComGD, anti-ComGE, anti-ComGF and anti-ComGG,). Following three 10-min washes with PBST, the membranes were incubated for 1 h with an anti-rabbit secondary antibody conjugated to horseradish peroxidase (GE Healthcare), diluted 1/10,000 in PBST. The membranes were washed again three times for 10 min in PBST, dried and developed with Amersham ECL Prime Western Blotting Detection Reagent (GE Healthcare) or SuperSignal West Atto Ultimate Sensivity Chemiluminescent Substrate (Thermo Scientific).

### Bioinformatics

DNA Strider [66] was used to routinely handle and analyse protein sequences. We identified protein domains by using Interposal [47]. We used AlphaFold [48] for modelling 3D structures Com proteins from *S. sanguinis* 2908. Pilins were modelled under their mature form, upon manually removing the N-terminal leader peptides. The AlphaFold parameters we usually applied consisted in generating five predictions in total with a final relaxation step included, after choosing the suitable model configuration (monomer or multimer). The predictions were ranked according to per-residue confidence metrics – on a scale from 0 to 100 – pLDDT (for monomers) or ipTM+pTM (for multimers). We always chose the prediction with the highest confidence score. We used PyMOL (Schrödinger) for visualising 3D structures and generating figures in this manuscript.

## Acknowledgements

This work was supported by the Agence Nationale de la Recherche (ANR-21-CE11-0008-01). We are grateful to Artemis Kosta and Hugo Le Guenno (Plateforme de Microscopie, Institut de Microbiologie de la Méditerranée, Marseille) for help with transmission electron microscopy. We would like to thank Sophie Helaine (Harvard Medical School), Emilia Mauriello (Laboratoire de Chimie Bactérienne) and Romé Voulhoux (Laboratoire de Chimie Bactérienne) for critical reading of this manuscript.

## Notes

### Competing Interest Statement

The authors have declared no competing interest.

