## Supplementary Information for "Systematic functional analysis of the Com pilus in *Streptococcus sanguinis*: a minimalistic type 4 filament dedicated to DNA uptake in monoderm bacteria"

### Table S1. Bacterial strains used in this study.

| Name | Description | Source |
|---|---|---|
| 2908 | naturally competent throat isolate | [1] |
| $Str^R$ 2908 | spontaneous mutant resistant to streptomycin (*$rpsL_{A167G}$*) | [1] |
| Δ*pil* Δ*fim* | Δ*pil* Δ*fim* | this study |
| $P_{ldh}$ *comX* | Δ*pil*::$P_{ldh}$ *comX-ermAM* | this study |
| Δ*comGA* | Δ*comGA*::*aphA3* | this study |
| Δ*comGA*::*comGA* | Δ*comGA*::*aphA3* Δ*pil*::$P_{ldh}$ *comGA-ermAM* | this study |
| Δ*comGB* | Δ*comGB*::*aphA3* | this study |
| Δ*comGB*::*comGB* | Δ*comGB*::*aphA3* Δ*pil*::$P_{ldh}$ *comGB-ermAM* | this study |
| Δ*comGC* | Δ*comGC*::*aphA3* | this study |
| Δ*comGC*::*comGC* | Δ*comGC*::*aphA3* Δ*pil*::$P_{ldh}$ *comGC-ermAM* | this study |
| Δ*comGD* | Δ*comGD*::*aphA3* | this study |
| Δ*comGD*::*comGD* | Δ*comGD*::*aphA3* Δ*pil*::$P_{ldh}$ *comGD-ermAM* | this study |
| Δ*comGE* | Δ*comGE*::*aphA3* | this study |
| Δ*comGE*::*comGE* | Δ*comGE*::*aphA3* Δ*pil*::$P_{ldh}$ *comGE-ermAM* | this study |
| Δ*comGF* | Δ*comGF*::*aphA3* | this study |
| Δ*comGF*::*comGE* | Δ*comGF*::*aphA3* Δ*pil*::$P_{ldh}$ *comGF-ermAM* | this study |
| Δ*comGG* | Δ*comGG*::*aphA3* | this study |
| Δ*comGG*::*comGG* | Δ*comGG*::*aphA3* Δ*pil*::$P_{ldh}$ *comGG-ermAM* | this study |
| Δ*comC* | Δ*comC*::*aphA3* | this study |
| Δ*comC*::*comC* | Δ*comC*::*aphA3* Δ*pil*::$P_{ldh}$ *comC-ermAM* | this study |
| $P_{ldh}$ *comG* | $P_{ldh}$ *comGA-GB-GC-GD-GE-GF-GG* | this study |
| $P_{ldh}$ *comG* $P_{ldh}$ *comC* | $P_{ldh}$ *comGA-GB-GC-GD-GE-GF-GG* Δ*pil*::$P_{ldh}$ *comC-ermAM* | this study |

Δ, deleted gene(s).
::, inserted gene(s).
*pil*, 22 kb cluster of genes encoding T4aP [1].
*fim*, 5.4 kb cluster of genes encoding sortase-assembled fimbriae [2].
*aphA3*, cassette encoding an aminoglycoside phosphotransferase conferring resistance to kanamycin.
*ermAM*, cassette encoding an rRNA methyltransferase conferring resistance to erythromycin.
$P_{ldh}$, promoter of the gene encoding the lactate dehydrogenase in 2908.

### Table S2. Primers used in this study.

| Name | Sequence* |
|---|---|
| **Construction of marked deletion mutants** | |
| *aphA3*-F | ATGGCTAAAATGAGAATATCACC |
| *aphA3*-R | CTAAAACAATTCATCCAGTAAAA |
| ΔGA-F1 | agccattgaagcagtaggga |
| ΔGA-R1 | GGTGATATTCTCATTTTAGCCATacatcctcctcaccttactat |
| ΔGA-F2 | TTTTACTGGATGAATTGTTTTAGGGGAGGAGGTGTCGTTGATT |
| ΔGA-R2 | agagcattgccgcataaagt |
| ΔGB-F1 | actgtgaggaagccaaggtt |
| ΔGB-R1 | GGTGATATTCTCATTTTAGCCATcttaggcgtagacaatttttc |
| ΔGB-F2 | TTTTACTGGATGAATTGTTTTAGTGTTTTACTTTATGCGGCAATGC |
| ΔGB-R2 | gtcctgtcctccgtctgaaa |
| ΔGC-F1 | tcttcagcagcggttttcac |
| ΔGC-R1 | GGTGATATTCTCATTTTAGCCATtataaatgaacctccatattctga |
| ΔGC-F2 | TTTTACTGGATGAATTGTTTTAGAGCTCAGGAAATATCAGCCAGA |
| ΔGC-R2 | aagtccctgccatcttgtct |
| ΔGD-F1 | ttatcatgctgggattgcgc |
| ΔGD-R1 | GGTGATATTCTCATTTTAGCCATcttaattggcaaccgtttgag |
| ΔGD-F2 | TTTTACTGGATGAATTGTTTTAGTTTCAGACGGAGGACAGGA |
| ΔGD-R2 | tgttgaaaagctgcaaacgc |
| ΔGE-F1 | aggagatcggagcagacatg |
| ΔGE-R1 | GGTGATATTCTCATTTTAGCCATtctagcttgaagctgtcgtc |
| ΔGE-F2 | TTTTACTGGATGAATTGTTTTAGCACGAAGGGGAGGTAGTCAT |
| ΔGE-R2 | agtccgacttaccagcttcg |
| ΔGF-F1 | cagcggtggtcaaggtagta |
| ΔGF-R1 | GGTGATATTCTCATTTTAGCCATtttaaggtttttgaacacggatg |
| ΔGF-F2 | TTTTACTGGATGAATTGTTTTAGAGGTTTAGAGAGGGAGTTCGTC |
| ΔGF-R2 | catcagcttcctctccgtga |
| ΔGG-F1 | atttcaggccggcagatttc |
| ΔGG-R1 | GGTGATATTCTCATTTTAGCCATctcaactttcttcttccacac |
| ΔGG-F2 | TTTTACTGGATGAATTGTTTTAGAGAGAAAGCGATTTTGGACGA |
| ΔGG-R2 | agtgtgctcatccggactag |
| ΔC-F1 | cctccgcctcaaaataagc |
| ΔC-R1 | GGTGATATTCTCATTTTAGCCATacttatttattcgtaattttaaacag |
| ΔC-F2 | TTTTACTGGATGAATTGTTTTAGCTGTCTTTTGTTTGGTGTGG |
| ΔC-R2 | cgttgaaagtagaggcgt |
| **Construction of unmarked deletion mutants** | |
| *pheS*-F | ATGACGAAAACGATTGAAGAAC |
| *aph*-R | CTAAAACAATTCATCCAGTAAAA |
| Δ*pil*-F1 | gagaaagcgacaaggaggtg |
| Δ*pil*-R1 | GTTCTTCAATCGTTTTCGTCATttgtttctcctgtctgtgatttt |
| Δ*pil*-F2 | TTTTACTGGATGAATTGTTTTAGCTTTGAACTCAGACAGAAAGGGG |
| Δ*pil*-R2 | tacacatgatccccagccag |
| Δ*pil*-R3 | CCCCTTTCTGTCTGAGTTCAAAGTTGTTTCTCCTGTCTGTGATTTT |
| Δ*pil*-F3 | AAAATCACAGACAGGAGAAACAACTTTGAACTCAGACAGAAAGGGG |
| Δ*fim*-F1 | gccaagcacctgactagtag |
| Δ*fim*-R1 | GCTCGGCTCTTTTCTTTTCCgtttggtcttcgaggcat |
| Δ*fim*-F2 | atgcctcgaagaccaaacGGAAAAGAAAAGAGCCGAGC |
| Δ*fim*-R2 | attccaccgcgtcatcaatg |
| **Complementation of deletion mutants** | |
| $P_{ldh}$ *comX-F1* | agccacgataacactcatag |
| $P_{ldh}$ *comX-R1* | ACCTTCTCATAAGTAGTTCTAAAATCCATttctaaacatctccttattttttagg |
| $P_{ldh}$ *comX-F2* | cctaaaaaaataaggagatgtttagaaATGGATTTTAGAACTACTTATGAGAAGGT |
| $P_{ldh}$ *comX-R2* | TTATAAAATCGTATTGTCTTGAAAATC |
| $P_{ldh}$ *comXerm-F1* | cccgagccacgataacactc |
| $P_{ldh}$ *comXerm-R1* | GTTCATtgataatatctcctTTATAAAATCGTATTGTCTTGAAAATC |
| $P_{ldh}$ *comXerm-F2* | TTATAAaggagatattatcaATGAACAAAAATATAAAATATTCTC |
| $P_{ldh}$ *comXerm-R2* | TTATTTCCTCCCGTTAAATAATAG |
| comp-F1 | tacacatgatccccagccag |
| compX-R1 | CTAtgagtgttatcgtggctcCTTTGAACTCAGACAGAAAGG |
| compX-F2 | AGCCACGATAACACTCATAG |
| compX-R2 | TTATTTCCTCCCGTTAAATAATAG |
| compX-F3 | CTATTATTTAACGGGAGGAAATAAttgtttctcctgtctgtgattt |
| comp-R3 | gagaaagcgacaaggaggtg |
| comp-F1 | tacacatgatccccagccag |
| compGA-R1 | GCAATTTCTTGAACCATttctaaacatctccttatttttttagg |
| compGA-F2 | ggagatgtttagaaATGGTTCAAGAAATTGCAAAAGAAATGATCAGG |
| compGA-R2 | GTTCATtgataatatctcctTTAGGCGTAGACAATTTTTTCGG |
| compGA-F3 | CCGAAAAAATTGTCTACGCCTAAaggagatattatcaATGAACA |

| | |
|---|---|
| comp-R3 | gagaaagcgacaaggaggtg |
| comp-F1 | tacacatgatccccagccag |
| compGB-R1 | CCTTTGCTCAGCACttctaaacatctccttatttttttagg |
| compGB-F2 | ggagatgtttagaaGTGCTGAGCAAAGGCAGACCGAAAAAATTGTC |
| compGB-R2 | GTTCATtgataatatctcctTTATAAATGAACCTCC |
| compGB-F3 | GGAGGTTCATTTATAAaggagatattatcaATGAACA |
| comp-R3 | gagaaagcgacaaggaggtg |
| comp-F1 | tacacatgatccccagccag |
| compGC-R1 | GTTAAGTTTTTCATttctaaacatctccttatttttttagg |
| compGC-F2 | ggagatgtttagaaATGAAAAACTTAACACCTTAAAAGTTCAAGCATTCACCC |
| compGC-R2 | GTTCATtgataatatctcctTTAATTGGCAACCGTTTGAGTT |
| compGC-F3 | CTCAAACGGTTGCCAATTAAaggagatattatcaATGAACAA |
| comp-R3 | gagaaagcgacaaggaggtg |
| comp-F1 | tacacatgatccccagccag |
| compGD-R1 | CCACTGTGTTTTCCATttctaaacatctccttatttttttagg |
| compGD-F2 | ggagatgtttagaaATGGAAAACACAGTGGCGAAACTCAAACGG |
| compGD-R2 | GTTCATtgataatatctcctCTAGCTTGAAGCTGTCGTCTTTTTAAACTTGCC |
| compGD-F3 | CGACAGCTTCAAGCTAGaggagatattatcaATGAACA |
| comp-R3 | gagaaagcgacaaggaggtg |
| comp-F1 | tacacatgatccccagccag |
| compGE-R1 | CGTCTTTTTAAACTTGCCATttctaaacatctccttatttttttagg |
| compGE-F2 | ggagatgtttagaaATGGCAAGTTTAAAAAGACGACAGC |
| compGE-R2 | GTTCATtgataatatctcctTTAAGGTTTTTGAACACGGATGACTACC |
| compGE-F3 | GGTAGTCATCCGTGTTCAAAAACCTTAAaggagatattatcaATGAACA |
| comp-R3 | gagaaagcgacaaggaggtg |
| comp-F1 | tacacatgatccccagccag |
| compGF-R1 | CCTTGACTTTAAGGTTTTTGAACACttctaaacatctccttatttttttagg |
| compGF-F2 | ggagatgtttagaaGTGTTCAAAAACCTTAAAGTCAAGG |
| compGF-R2 | GTTCATtgataatatctcctTCAACTTTCTTCTTCCACACG |
| compGF-F3 | GGAAGAAGAAAGTTGAaggagatattatcaATGAACA |
| comp-R3 | gagaaagcgacaaggaggtg |
| comp-F1 | tacacatgatccccagccag |
| compGG-R1 | CCTCAACTTTCTTCTTCCACACttctaaacatctccttatttttttagg |
| compGG-F2 | ggagatgtttagaaGTGTGGAAGAAGAAAGTTGAGG |
| compGG-R2 | GTTCATtgataatatctcctTTAGTCCGACTTACCAGCTTCGTCC |
| compGG-F3 | GGACGAAGCTGGTAAGTCGGACTAAaggagatattatcaATGAACA |
| comp-R3 | gagaaagcgacaaggaggtg |
| comp-F1 | tacacatgatccccagccag |
| compC-R1 | GTATAGATGAATCATttctaaacatctccttatttttttagg |
| compC-F2 | ggagatgtttagaaATGATTCATCTATACTTTTTTCTCC |
| compC-R2 | GTTCATtgataatatctcctTCAGTAAAAGAGTAGGG |
| compC-F3 | CCCTACTCTTTTACTGAaggagatattatcaATGAACA |
| comp-R3 | gagaaagcgacaaggaggtg |
| **Construction of a strain expressing Com pili constitutively** | |
| $P_{ldh}$ *GA*-F1 | CTCCGTCAAAGATTATCATGG |
| $P_{ldh}$ *GA*-R1 | gatgtaaacgtttttacaaatggTTAAAGAGGCTTAGCGGTTTTT |
| $P_{ldh}$ *GA*-F2 | CTAAAAACCGCTAAGCCTCTTTAAccatttgtaaaaacgtttacatc |
| $P_{ldh}$ *GA*-R2 | TTAGGCGTAGACAATTTTTTCGG |
| **Generating a PCR product for quantifying competence** | |
| *rpsL*-F | ggcaggtgtagctgtctttg |
| r*psL*-R | ctcttgctccatccagtcca |

*Genes are in upper case; flanking regions are in lower case. Regions of complementarity for splicing PCR are underlined.

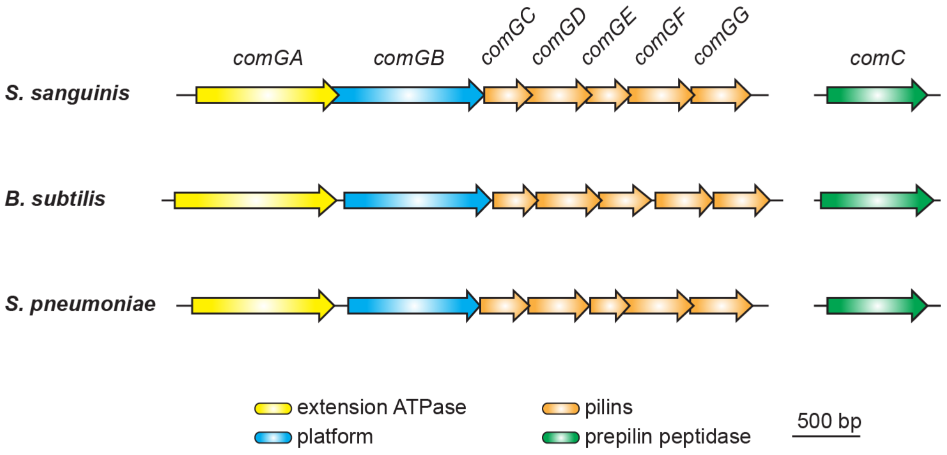


**Fig. S1. Genomic organisation of the genes involved in the synthesis of the Com pilus in model competent species.** The genes, with the same colour code as in Fig. 3, are drawn to scale. The corresponding proteins are listed at the bottom. The chosen strains are *S. sanguinis* 2908, *B. subtilis* 168 and *S. pneumoniae* TIGR4.

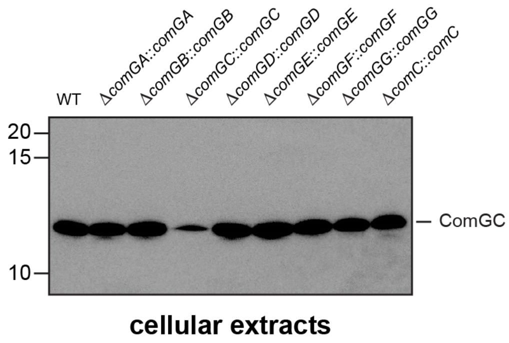


**Fig. S2. ComGC in deletion mutants in the eight *com* genes complemented with a WT copy of the corresponding genes.** Immunoblotting was performed on cellular extracts using an anti-ComGC antibody. The WT strain is included as a control. Extracts were prepared from cultures equalised to the same $OD_{600}$, and equivalent volumes were loaded in each lane. Molecular weight markers (in kDa) are indicated on the left.

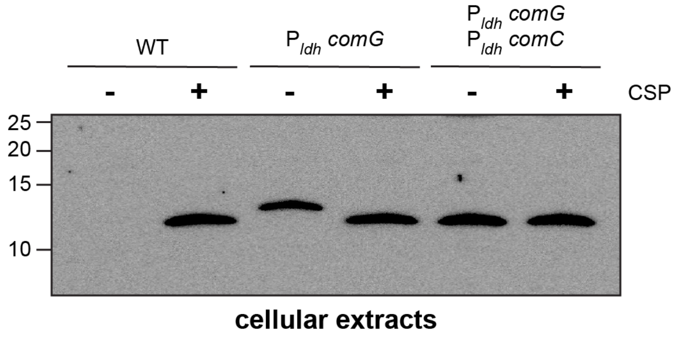


**Fig. S3. Detection of ComGC by immunoblotting in a strain expressing Com pili constitutively.** Cellular extracts were prepared +/- CSP induction from cultures of WT, $P_{ldh}$ *comG* (intermediate strain), and $P_{ldh}$ *comG* $P_{ldh}$ *comC* (final strain) adjusted to the same $OD_{600}$. Immunoblotting was performed using an anti-ComGC antibody. Molecular weight markers (in kDa) are indicated on the left.

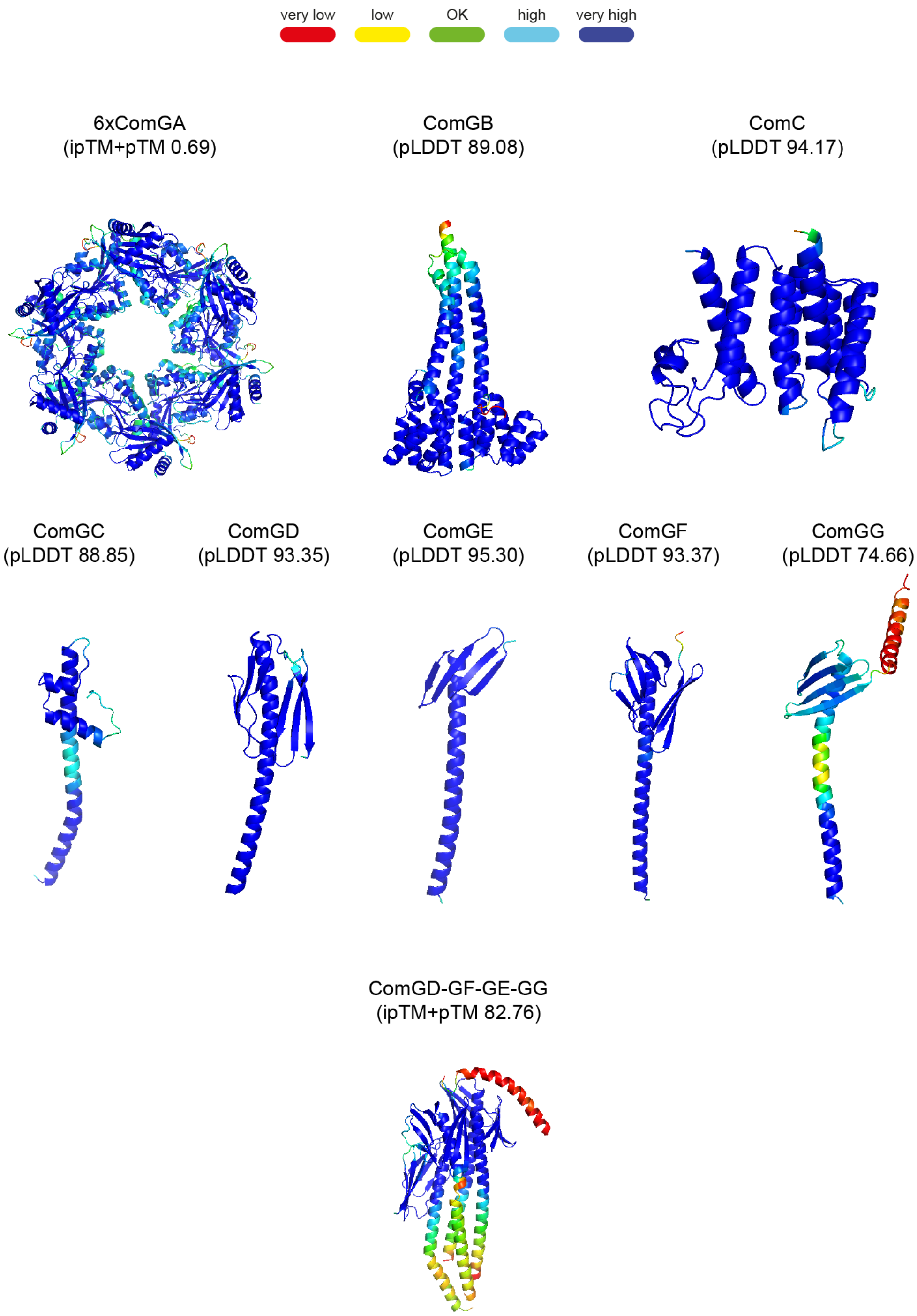


**Fig. S4. Accuracy of the AlphaFold models.** The per-residue confidence scores given by AlphaFold [3] – pLDDT (monomers) or ipTM+pTM (multimers) ranging from 0 to 100 – are indicated for each model. The structures are coloured by confidence measures from dark blue (very high accuracy expected) to red (very low accuracy, which should not be interpreted and may be disordered).

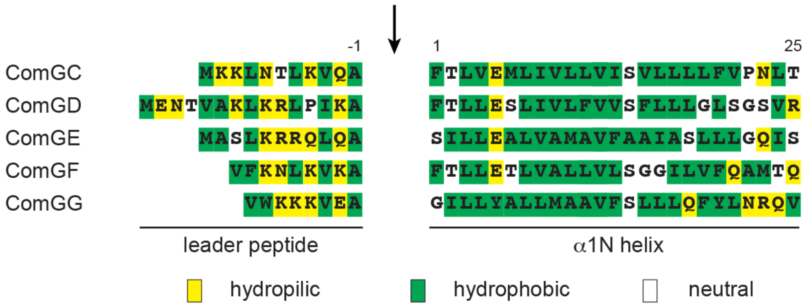


**Fig. S5. Sequence alignment of the N-terminal class 3 signal peptides of the five Com pilins in *S. sanguinis* 2908.** The 8-15 residues long leader peptides, which end with a conserved Ala, contain a majority of hydrophilic (shaded in yellow) or neutral (no shading) residues. The leader peptides are cleaved (indicated by the vertical arrow) by the PPase ComC. The mature proteins start with a tract of 21 predominantly hydrophobic residues (shaded in green), which invariably form the protruding N-terminal half of an extended α-helix (α1N). α1N is the main assembly interface of pilins within filaments.

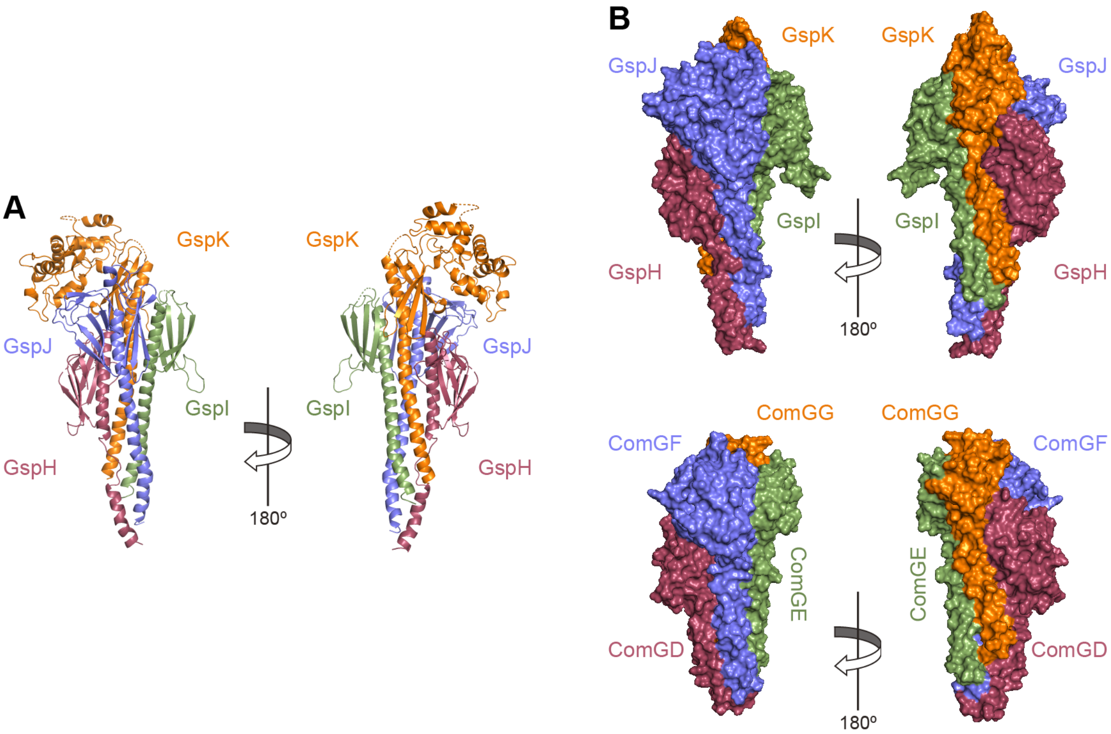


**Fig. S6. Structural similarity between tip-located complexes of four minor pilins in Com pili and T2SS.** The structures – 180° views – are shown as cartoon or surface. Homologous subunits in the two T4F are highlighted with the same colour (we used the same colour code as in Fig. 10). **A**) The GspH-GspJ-GspI-GspK complex from *Pseudomonas aeruginosa* T2SS [4]. **B**) Side-by side comparison of the H-J-I-K and D-F-E-G complexes. To illustrate the structural similarity, we have removed the disordered C-terminus of ComGG and the extra module in GspK.
